# The miR-221-5p/RAD18/RAD51 Axis Regulates DNA Damage Tolerance and Homologous Recombination to Drive Platinum Resistance in Ovarian Cancer

**DOI:** 10.64898/2026.05.11.724004

**Authors:** Tasmin Rahman Omy, Naresh Sah, Subash Kairamkonda, Chinnadurai Mani, Md Ariful Islam, Mark B. Reedy, Komaraiah Palle

## Abstract

Platinum resistance remains a major barrier in Ovarian cancer (OC) treatment[1]. While hyperactivation of DNA damage response (DDR) is a hallmark of chemoresistance[2], the underlying epigenetic mechanisms driving this adaptation remain poorly understood. Here, we identify a novel post-transcriptional regulatory axis involving miR-221-5p that governs two critical DDR effectors: RAD18, which mediates DNA damage tolerance through trans-lesion synthesis (TLS)[3][4], and RAD51, the central recombinase for homologous recombination (HR)[5][6]. Although the miR-221/222 cluster is traditionally categorized as oncogenic[7][8], we demonstrate that the miR-221-5p arm functions as a potent tumor suppressor in OC. Bioinformatic and luciferase reporter assays confirmed that miR-221-5p directly targets the 3′UTRs of both *RAD18* and *RAD51*. In OC clinical specimens and cell lines, miR-221-5p downregulation inversely correlates with RAD18/RAD51 expression. Functionally, miR-221-5p restoration suppressed platinum-induced PCNA mono-ubiquitination and HR, inducing a “functional BRCAness” that sensitized both established and patient-derived primary OC cells to carboplatin and PARP inhibition. Furthermore, *in vivo* disseminated xenograft models demonstrated that stable miR-221-5p expression significantly reduced tumor burden. Collectively, our results delineate a novel regulatory mechanism where loss of miR-221-5p drives chemoresistance by derepressing the RAD18/RAD51 axis, identifying this axis as a promising therapeutic target.

## Introduction

Ovarian cancer (OC) remains the most lethal gynecological malignancy worldwide[9], characterized by insidious progression and often diagnosed at an advanced stage. Although radical debulking surgery followed by platinum-taxane chemotherapy initially achieves high response rates, approximately 70% of patients succumb to disease recurrence. This clinical relapse is almost universally driven by acquired chemoresistance, which represents the most formidable barrier to long-term survival. Projections for 2025 underscore this persistent burden, with over 20,000 new cases and 12,000 deaths estimated in the U.S. alone [10].

The development of chemoresistance is a multifactorial process, yet the hyperactivation of the DNA damage response (DDR) is a hallmark strategy for tumor cell survival[11]–[13]. By efficiently repairing or bypassing the DNA crosslinks and double-strand breaks induced by platinum agents, cancer cells maintain genomic integrity and evade apoptosis[14][15]. While genetic alterations are primary drivers of DDR dysregulation, emerging evidence highlights epigenetic reprogramming, specifically the loss of regulatory microRNAs (miRNAs) as a critical posttranscriptional mechanism driving DDR adaptation[16][17].

Among these regulators, the miR-221/222 cluster is widely recognized for its oncogenic potential[18][19][20][21]. However, most research has focused on the 3p arm, leaving the specific functional role of the miR-221-5p strand in OC largely unexplored. In this study, we identify miR-221-5p as a master regulator of two critical, yet functionally distinct, DDR effectors: RAD18 and RAD51. These proteins constitute a dual-arm survival mechanism; RAD18 orchestrates DNA damage tolerance (DDT) via translesion synthesis (TLS), allowing cells to replicate past bulky platinum adducts[22], while RAD51 is the central recombinase for high-fidelity homologous recombination (HR) repair of DSBs[23]. The simultaneous upregulation of these factors in platinum-resistant OC suggests a coordinated network that shields the cancer genome from diverse therapeutic insults.

Using bioinformatic screening, we identified conserved miR-221-5p binding sites within the 3′ untranslated regions (3′-UTR) of both *RAD18* and *RAD51*[24][25]. We hypothesized that the downregulation or loss of miR-221-5p serves as a derepressing mechanism for the RAD18/RAD51 axis, thereby driving the increased expression of these repair proteins to promote DNA damage adaptation in OC cells. Using a combination of luciferase reporter assays, 3D organoid cultures, and *in vivo* xenograft models, we demonstrate that restoring miR-221-5p effectively collapses this repair axis, inducing a state of “functional BRCAness”. This restoration not only induces homologous recombination deficiency (HRD) and increases persistent DNA strand breaks, but also significantly re-sensitizes resistant OC cells to carboplatin and PARP inhibitor, olaparib. Collectively, our findings delineate a novel miR-221-5p/RAD18/RAD51 epigenetic regulatory axis and propose miR-221-5p replacement as a precision oncology strategy to overcome the formidable challenge of chemoresistance in aggressive ovarian cancer.

## Results

### RAD18 and RAD51 Overexpression Correlates with Poor Prognosis and miR-221-5p Downregulation in Ovarian Cancer

To investigate the clinical relevance of DNA damage response (DDR) hyperactivation in ovarian cancer (OC), we first interrogated the prognostic significance of two central DDR effectors: RAD51, the primary recombinase facilitating homologous recombination (HR), and RAD18, the E3 ubiquitin ligase essential for translesion synthesis (TLS). Analysis of The Cancer Genome Atlas (TCGA) datasets via GEPIA [26] and KMplotter [27] revealed that high mRNA expression of both *RAD51* (Fig. 1A) and *RAD18* (Fig. 1B) significantly correlates with reduced overall survival (OS) and progression-free survival (PFS) in OC patients (Fig. 1C, 1D). Validation at the protein level across a panel of OC cell lines confirmed a robust upregulation of RAD51 and RAD18 compared to non-malignant fallopian tube epithelial (FTEC) cells (Fig. 1E, 1F).

**Figure 1.**
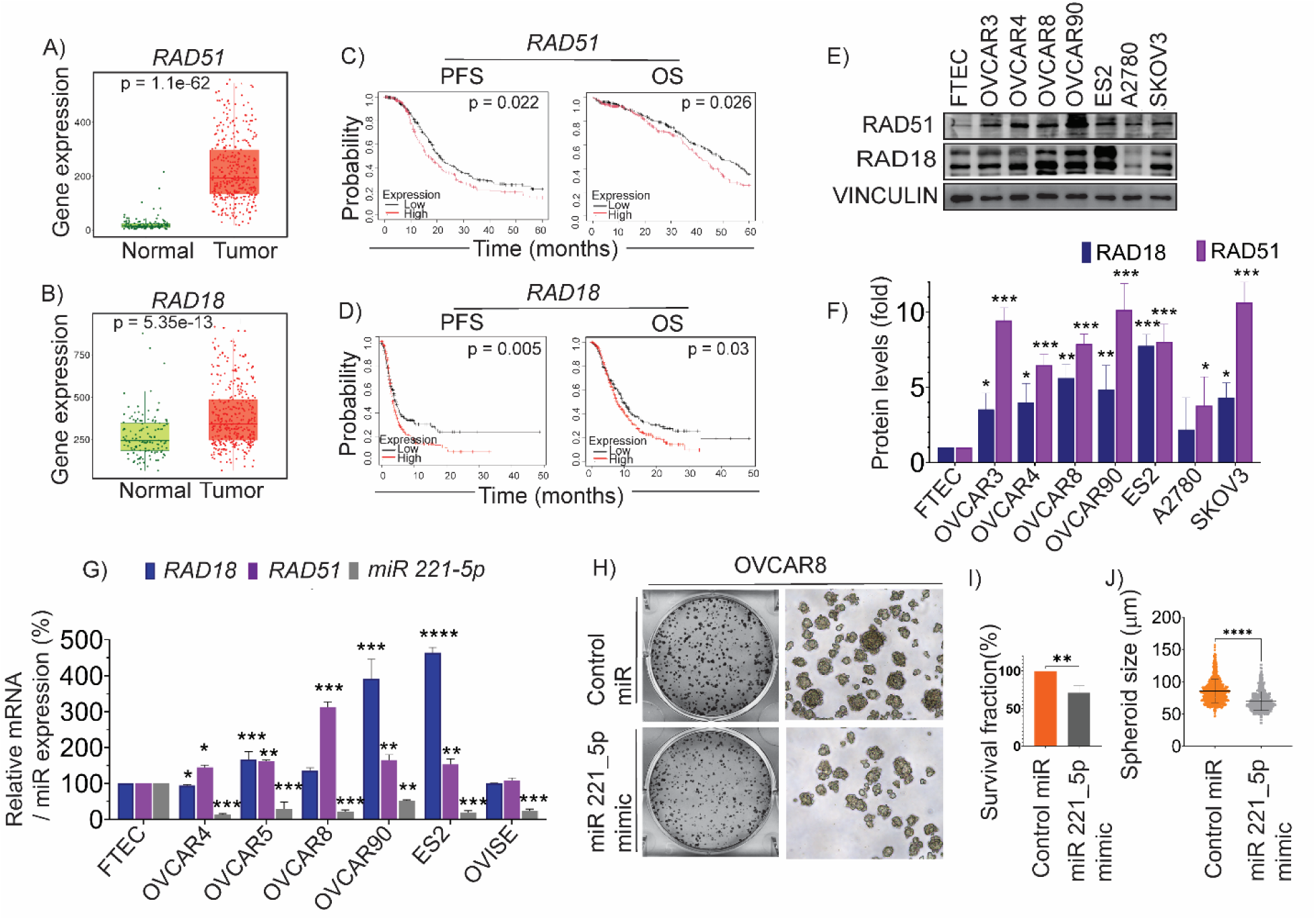
RAD18 and RAD51 overexpression correlates with poor prognosis and is regulated by miR-221-5p in ovarian cancer. (A, **B)** Box plots summarizing mRNA levels of *RAD51* (A) and *RAD18* (B) in the TCGA ovarian serous cystadenocarcinoma cohort. Statistical significance determined by two-tailed Student’s - test. **(C, D)** Kaplan–Meier survival curves for progression-free survival (PFS) and overall survival (OS) in patients stratified by high vs. low transcript expression of *RAD51* (C) or *RAD18* (D). values were calculated using the log-rank test. **(E, F)** Western blot analysis (E) and corresponding densitometric quantification (F) of RAD51 and RAD18 in non-malignant fallopian tube epithelial cells (FTEC) and a panel of ovarian cancer cell lines. Vinculin served as the loading control. Data represent the mean ± SD of ≥3 independent experiments (*P*<0.05, \**P*<0.01, \*\**P*<0.001; two-tailed *t*-test). (G) Relative expressions of *RAD18*, *RAD51*, and miR-221-5p in FTEC and OC cell lines as determined by qRT-PCR. mRNA and miRNA levels were normalized to *GAPDH* and *U6*, respectively, and plotted as a percentage of FTEC expression. (Values represent mean ± SD from ≥3 independent experiments (*P*<0.05, \**P*<0.01, \*\**P*<0.001; two-tailed *t*-test). (H) Representative images of low-density colony assays (left) and 3D spheroid cultures (right) in OVCAR8 cells following transfection with control miRNA or miR-221-5p mimics. (I) Survival fraction from clonogenic assays in (H); bars represent mean ± SD from three biological replicates (\**P*<0.01; two-tailed *t*-test). (J) Violin plot of spheroid diameters measured from 3D assay in (H). . Each point represents an individual spheroid (n ≥100 spheroids per group, diameter larger than 50 μm). Data presented as mean ± SD; (*P*<0.05, \**P*<0.01, \*\**P*<0.001; two-tailed *t*-test).

Based on our previous findings regarding miRNA-mediated DDR regulation, we performed a re-analysis of the GEO dataset (GSE235525)[28]. This identified miR-221-5p as one of the most significantly downregulated miRNAs in clinical OC specimens compared to normal tissue (Supplementary Figure. 1A, 1B and Supplemental Table 1). To further characterize this regulatory network between *miR-221-5p* and *RAD51* and *RAD18*, we analyzed the expression profile in a panel of established OC cell lines and non-malignant FTEC controls. Quantitative RT-PCR (qRT-PCR) across our cell line panel established a striking inverse correlation. *miR-221-5p* was nearly undetectable in malignant cell lines where *RAD18* and *RAD51* were substantially overexpressed (Fig. 1G). Functionally, ectopic restoration of miR-221-5p in OVCAR8 cells significantly suppressed the malignant phenotype, as evidenced by a significant reduction in clonogenic survival and 3D spheroid volume (Fig. 1H-1J). Similar results were observed in ES2 (Supplementary Figure 1B, 1C) and SKOV3 cells (Supplementary Figure 1D-1G). Importantly, flow cytometric analysis indicated that these effects were not due to a general arrest in the cell cycle, as G1/S/G2 distributions remained stable (Supplementary Figure 4A-4D), suggesting that miR-221-5p acts primarily by impairing proliferative signaling and antitumorigenic effect.

### miR-221-5p Directly Targets the 3′UTRs of *RAD18* and *RAD51* to Suppress mRNA Expression

We next sought to determine whether the observed inverse correlation was driven by direct post-transcriptional silencing. Transfection of OVCAR8, ES2, and SKOV3 cells with miR-221-5p mimics resulted in a significant reduction in both RAD18 and RAD51 protein levels (Fig. 2A-2F). This downregulation was mirrored by a significant reduction in OVCAR8, ES2 respective mRNA transcripts (Supplementary Figure 2A, 2B), suggesting that miR-221-5p facilitates both mRNA degradation and translational inhibition.

**Figure 2.**
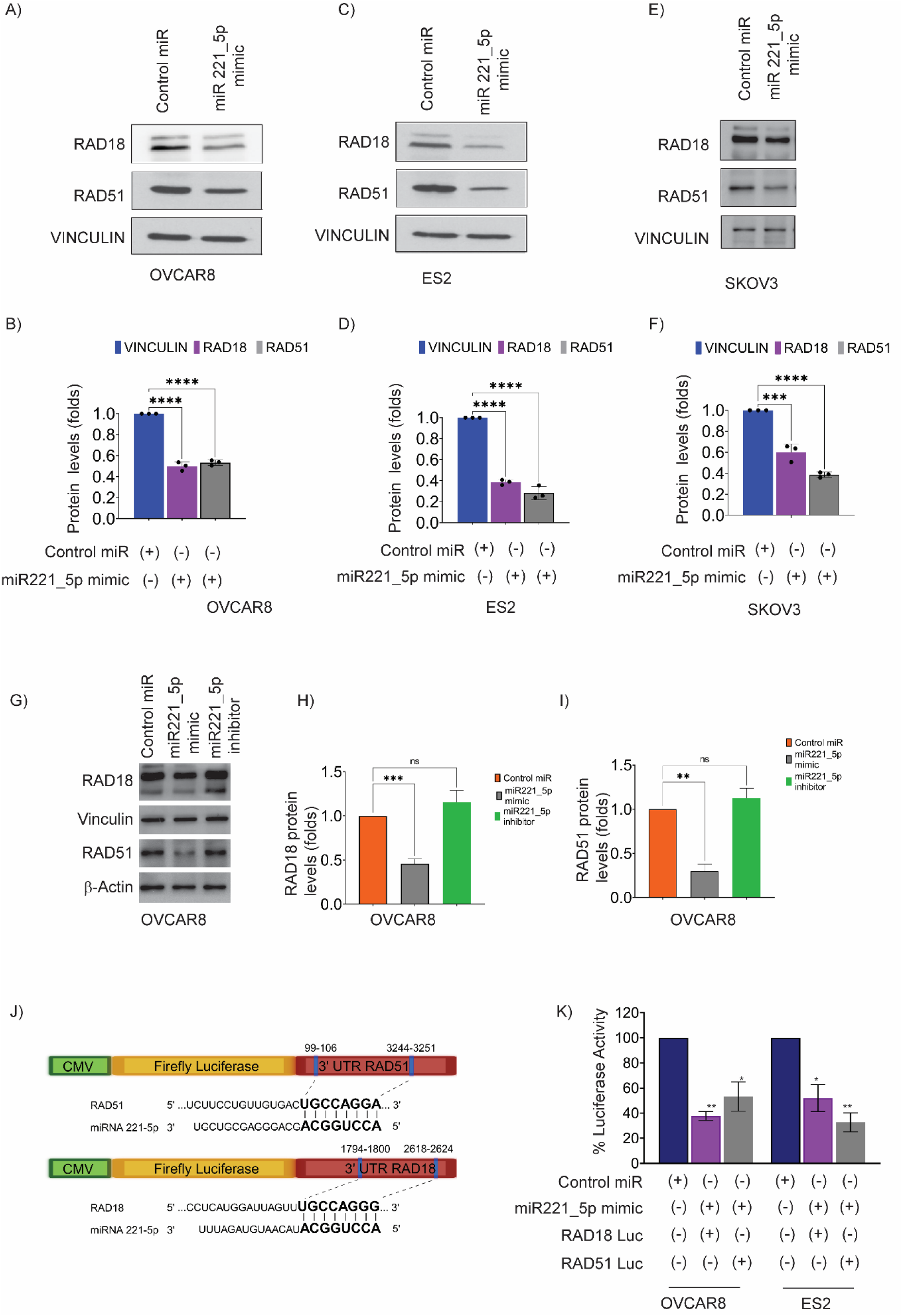
miR–221–5p directly suppresses RAD18 and RAD51 by targeting their 3′ UTRs. (A–F) Western blot analyses of RAD18 and RAD51 in OVCAR8 (A,B), ES2 (C,D) and SKOV3 (E,F) cells 48h after transfection with a control miR or a miR–221–5p mimic. Vinculin serves as loading control. Corresponding bar graphs in B, D and F show densitometric quantification of protein expression normalized to vinculin (mean ± SD, n = 3 independent replicates; ****P < 0.0001, two–tailed *t*-test). (G–I) Functional rescue experiment in OVCAR8 cells. Cells were transfected with control miR, miR–221–5p mimic or a miR–221–5p inhibitor (anti-miR). Immunoblot blot (G) and quantification (H, I) demonstrate that miR–221–5p-mediate downregulation of RAD18 and RAD51 effectively reversed by the inhibitor, confirming the specificity of this regulatory axis. (J) Schematic representation of firefly luciferase reporter constructs containing the 3′–untranslated regions (3′ UTRs) of *RAD51* or *RAD18*. Conserved miR–221–5p seed sequences within each 3′ UTR are highlighted. (K) Relative luciferase activity in OVCAR8 and ES2 cells co-transfected with miR-221-5p mimics and the indicated 3′ UTR reporter vectors. Values were normalized to internal controls. Data are presented as mean ± SD of n = 3; *P < 0.05, **P < 0.01, ***P < 0.001 by two–tailed t-test).

To further validate this regulatory axis, we performed a reciprocal rescue experiment using a miR-221-5p-specific inhibitor (anti-miR). While miR-221-5p mimics significantly suppressed RAD18 and RAD51 protein expression in OVCAR8 cells, the addition of the anti-miR inhibitor effectively rescued and replenished protein levels to baseline (Fig. 2G–2I). This restoration confirms that the downregulation of these repair factors is specifically mediated by miR-221-5p.

To provide molecular evidence for this interaction, we utilized *in silico* prediction algorithms (TargetScan and mirBase)[29][25][16][30], which identified highly conserved miR-221-5p seed sequences within the 3′UTRs of *RAD18* and *RAD51* (Fig. 2J). Reporter constructs were generated by fusing the luciferase open reading frame (ORF) to the 3′UTR regions of *RAD18* and *RAD51*, specifically containing the predicted miR-221-5p binding sites (nucleotides 1522–1528 and 1702–1708, respectively) (Supplementary Figure 2B). To confirm direct regulation, we performed dual-luciferase reporter assays for both the targets. In OVCAR8 and ES2 cell lines, ectopic expression of miR-221-5p mimics significantly reduced luciferase activity compared to non-targeting controls (Fig. 2K). Collectively, these findings provide definitive evidence that miR-221-5p acts as a direct negative regulator of the *RAD18/RAD51* axis.

### miR-221-5p Restoration Collapses TLS and HR Signaling and Sensitizes OC Cells to Carboplatin

To evaluate the functional consequences of miR-221-5p-mediated repression, we examined downstream DDR signaling. A critical step in translesion synthesis (TLS) is the RAD18-dependent mono-ubiquitination of PCNA (Ub-PCNA)[31][32], a modification that permits replication forks to bypass platinum-DNA adducts. Western blot analysis revealed that miR-221-5p restoration dramatically attenuated Ub-PCNA levels following carboplatin exposure across multiple OC cell lines (Fig. 3A–C). Similarly, we also observed a concomitant reduction in Ub-FANCD2, a central marker of the Fanconi Anemia (FA) pathway[33], suggesting miR-221-5p triggers a broader collapse of DNA damage tolerance mechanisms.

**Figure 3.**
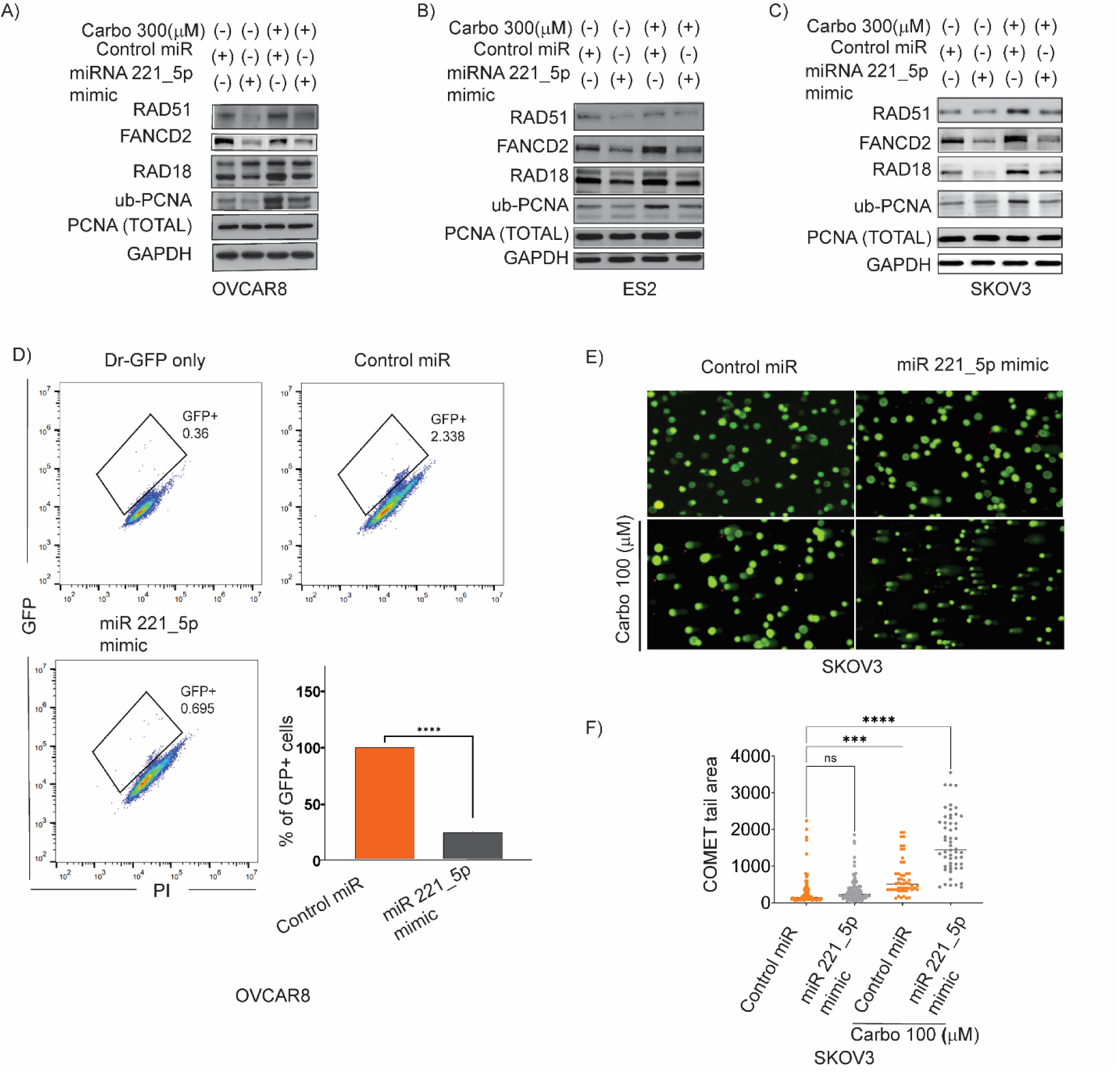
miR-221-5p restoration collapses DNA damage tolerance and homologous recombination, resensitizing carboplatin-induced damage. (A–C) Western blot analysis of OVCAR8 (A), ES2 (B) and SKOV3 (C) cells were transfected with a control miR or a miR-221-5p mimic and exposed to carboplatin (Carbo, 300 μM) for 24h. Total PCNA and GAPDH serve as loading controls. (D) OVCAR8 cells were transfected with pDR-GFP and selected using 5 µg/ml puromycin. Stably expressing cells were simultaneously transfected with miR-control or miR-221-5p mimics along with pCBASceI expression vector. After 48 h post-transfection, the GFP-positive cells was quantified via flow cytometry (BD Accuri) to determine the relative HR efficiency (mean ± SD of three independent experiments; ****P < 0.0001; two-tailed *t*-test). (E) Alkaline comet assay images of SKOV3 cells transfected with control miR or miR-221-5p mimic and treated with carboplatin (100 μM) for vehicle for 24h or vehicle. (F) Quantification of comet tail area from (E). Each point represents an individual nucleus; horizontal bars denote the mean ± SD (more than 25 cells per condition). (***P < 0.001, ****P < 0.0001; two-tailed *t*-test; ns, not significant).

We further quantified homologous recombination (HR) efficiency using DR-GFP reporter assays [33]in OVCAR8 cells. Ectopic miR-221-5p expression led to a significant decrease in GFP-positive (HR-proficient) cells following I-SceI-induced DNA double strand breaks (DSBs) (Fig. 3D). This repair deficiency was corroborated by alkaline comet assays, which demonstrated significantly elongated comet tails in miR-221-5p transfected cells, signifying the persistence of unresolved DNA strand breaks (Fig. 3E–3F). Immunofluorescence microscopy further revealed that while carboplatin-treated control cells formed robust RAD51 nuclear repair foci, miR-221-5p-treated cells failed to recruit RAD51 to damage sites. This was accompanied by a reciprocal increase in accumulation of 53BP1 foci, a marker of unrepaired DSBs (Fig. 4A–4D).

**Figure 4.**
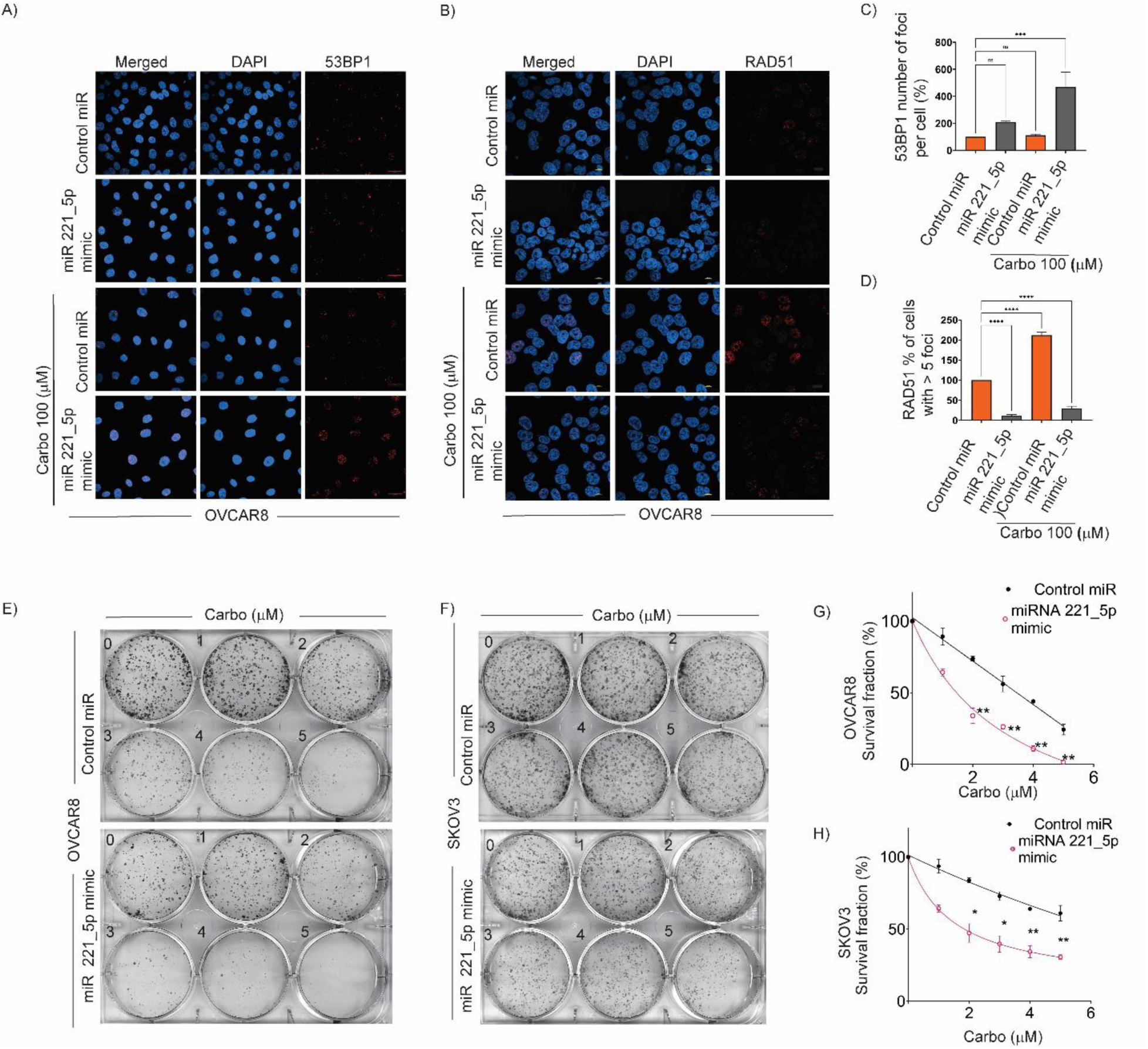
miR-221-5p disrupts RAD51 foci formation, promotes DNA damage accumulation, and sensitizes OC cells to carboplatin. (A,B) Representative immunofluorescence images of OVCAR8 cells transfected with control miRNA or miR-221-5p mimics, followed by treatment with vehicle or carboplatin (100 μM, 24 h). Cells were stained for 53BP1 (A), a marker of double-strand breaks, orRAD51 (B), a marker of homologous recombination repair. Scale bar represents 10 μm (C) Quantitative analysis of the percentage of cells positive for 53BP1 foci (≥ 5 foci/nucleus). Data represents the mean ± SEM from more than 75 cells per condition(*P < 0.05, **P < 0.01, ***P < 0.001 ; two-tailed *t*-test). (D) Quantitative analysis of the percentage of cells positive for RAD51 foci (≥ 5 foci/nucleus). Data represents the mean ± SEM from more than 75 cells per condition. (****P < 0.0001 ; two-tailed *t*-test). (E,F) Representative images of low-density colony assays using OVCAR8 (E) and SKOV3 (F) cells transfected with control or miR-221-5p mimics and treated with increasing doses of carboplatin (0–5 μM). (G,H) Clonogenic survival curves representing the survival fraction of OVCAR8 (G) and SKOV3 (H) cells from (E, F). Results illustrate significant chemosensitization following miR-221-5p restoration. Data represents the mean ± SEM of three independent experiments (**P < 0.01, ***P < 0.001 vs. control; two-tailed *t*-test).

To determine whether these molecular defects translate into enhanced therapeutic vulnerability, we performed colony formation assays in OVCAR8 and SKOV3 cells exposed to increased concentrations of carboplatin (up to 5 μM). While control-miR transfected cells exhibited a typical dose-dependent decrease in clonogenic survival, miR-221-5p restoration resulted significantly sensitized OC cells, resulting in a more pronounced reduction in colony forming capacity at all tested doses (Fig. 4E–4H). Notably, cell cycle profiling by flow cytometry revealed no significant redistribution in G1, S, or G2/M phases between control and miR-221-5p transfected cells (Supplementary Figure 4A-4D). This indicates that the observed carboplatin sensitization effect of miR-221-5p is not due to altered cell cycle kinetics, but rather a direct consequence of the dual impairment of RAD18-mediated TLS and RAD51-mediated HR.

### miR-221-5p Suppresses Growth, Stemness and Invasion in Patient-Derived Primary OC Models

To establish the clinical translatability of the miR-221-5p/RAD18/RAD51 axis, we extended our investigation to primary patient-derived OC cells (OV-TX-186 and OV-TX-285). Consistent with our findings in established cell lines, ectopic restoration of miR-221-5p effectively depleted both RAD18 and RAD51 protein levels in these primary models (Supplementary Figure 5A).

To assess the impact of this axis on the tumor-initiating capacity of primary cells, we utilized 3D organoid formation assays. miR-221-5p overexpression significantly compromised the organoid-forming efficiency and growth of both OV-TX-186 and OV-TX-285 cells (Fig. 5A–5D). This marked reduction in anchorage-independent growth reflects a loss of self-renewal capacity and stemness following miR-221-5p restoration. Furthermore, *in situ* immunofluorescence (IF) staining of the complex 3D architecture of OV-TX-186 organoids confirmed a robust downregulation of the target repair proteins, RAD18 and RAD51, within the organoid mass (Fig. 5E–5F).

**Figure 5.**
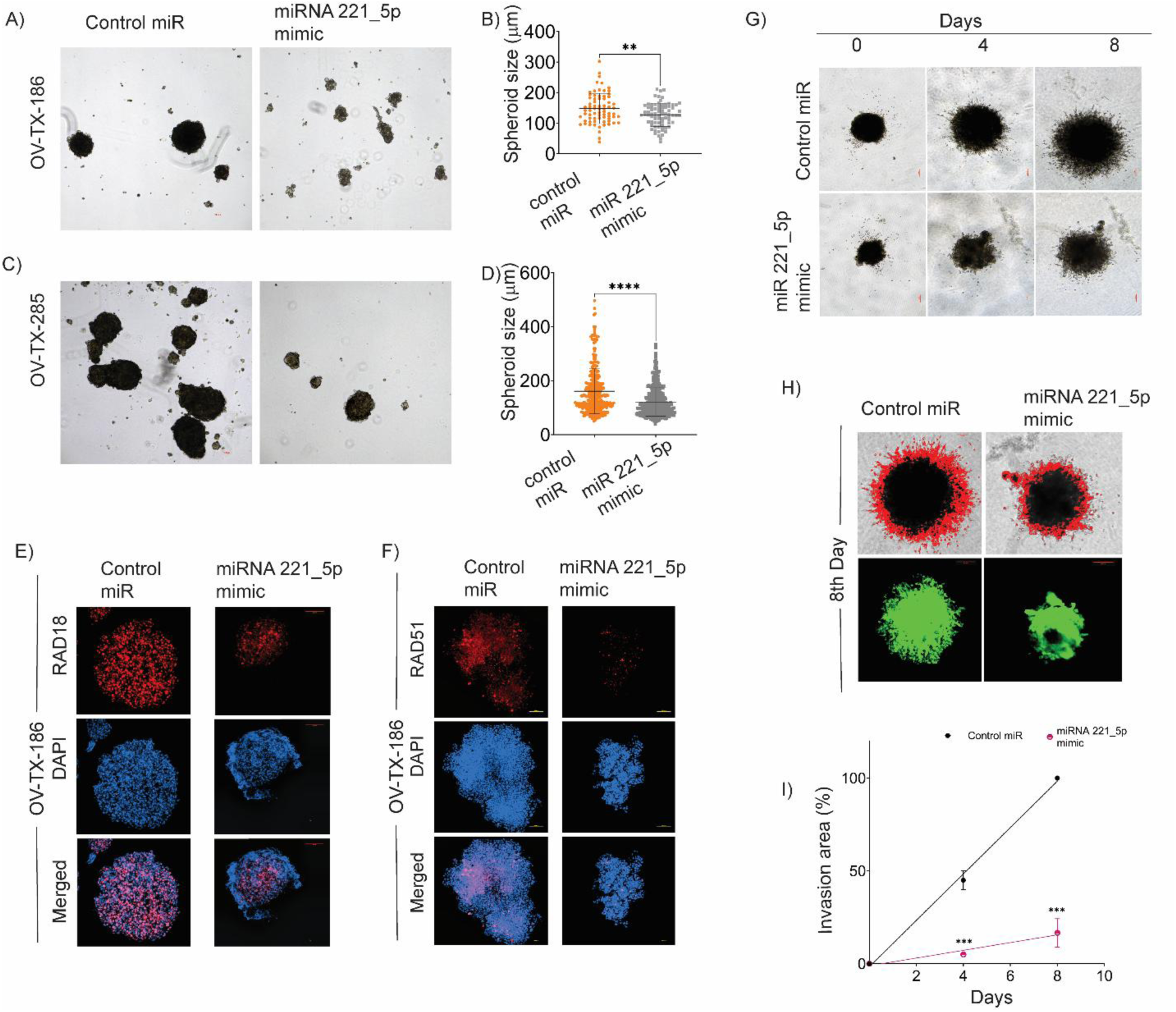
miR–221–5p attenuates proliferative capacity ss and invasion potential in primary patient–derived OC organoids. (A–D) Representative brightfield images (A, C) and quantitative size distribution violin plots (B, D) of 3D organoids derived from primary patient-derived OC models OV-TX-186 (A, B) and OV-TX-285 (C, D) post-transfection with control miRNA or miR-221-5p mimics. Each point in the violin plots represents an individual organoid (n ≥ 100 per condition with size greater than 50 μm; **P < 0.01, ****P < 0.0001; two-tailed *t*-test;). (E, F) Representative 3D confocal immunofluorescence images of OV-TX-186 organoids stained for RAD18 (E) or RAD51 (F) (red) following miR-221-5p restoration. Nuclei were counterstained with DAPI (blue).Scale bars, 50 μm. (G) Representative images of the 3D Matrigel-dependent invasion assay. OV-TX-186 organoids were embedded in a basement membrane matrix and monitored for invasive outward growth on days 0, 4, and 8. (H) High-magnification confocal images of day-8 organoids from (G) stained with fluorescent phalloidin (green) to visualize cytoskeletal reorganization and invadopodia formation during invasion. (I) Quantification of the relative invasion area (%) over an 8-day period (G). Scale bar represents 100 μm (n=3 per group; mean ± SD ; **P < 0.01, ***P < 0.001; two-tailed *t*-test).

Beyond self-renewal, we evaluated the role of miR-221-5p in modulating the metastatic potential of primary OC cells using 3D Matrigel invasion assays, which more accurately recapitulate the dynamic tumor microenvironment. Primary cells transfected with miR-221-5p mimics exhibited a significant attenuation in invasive potential compared to control-transfected cells (Fig. 5G–5I). Collectively, these data demonstrate that the miR-221-5p/RAD18/RAD51 axis is a critical regulator of OC stemness and invasiveness, further supporting the role of miR-221-5p as a potent suppressor of tumorigenesis in a clinically relevant context.

### miR-221-5p Overexpression Attenuates Ovarian Tumor Growth in a Disseminated Xenograft Model

To evaluate the therapeutic potential of miR-221-5p *in vivo*, SKOV3-Luc cells were stably transduced with lentiviral particles encoding either a miR-221-5p mimic or a non-targeting control. Stable integration and functional knockdown of the target proteins, RAD18 and RAD51, were confirmed via Western blot prior to implantation (Supplementary Fig. 6A). Furthermore, stable miR-221-5p expression significantly impaired *in vitro* 3D spheroid growth, validating the persistent tumor-suppressive effect of the lentiviral construct (Supplementary Fig. 6B–6C).

**Figure 6.**
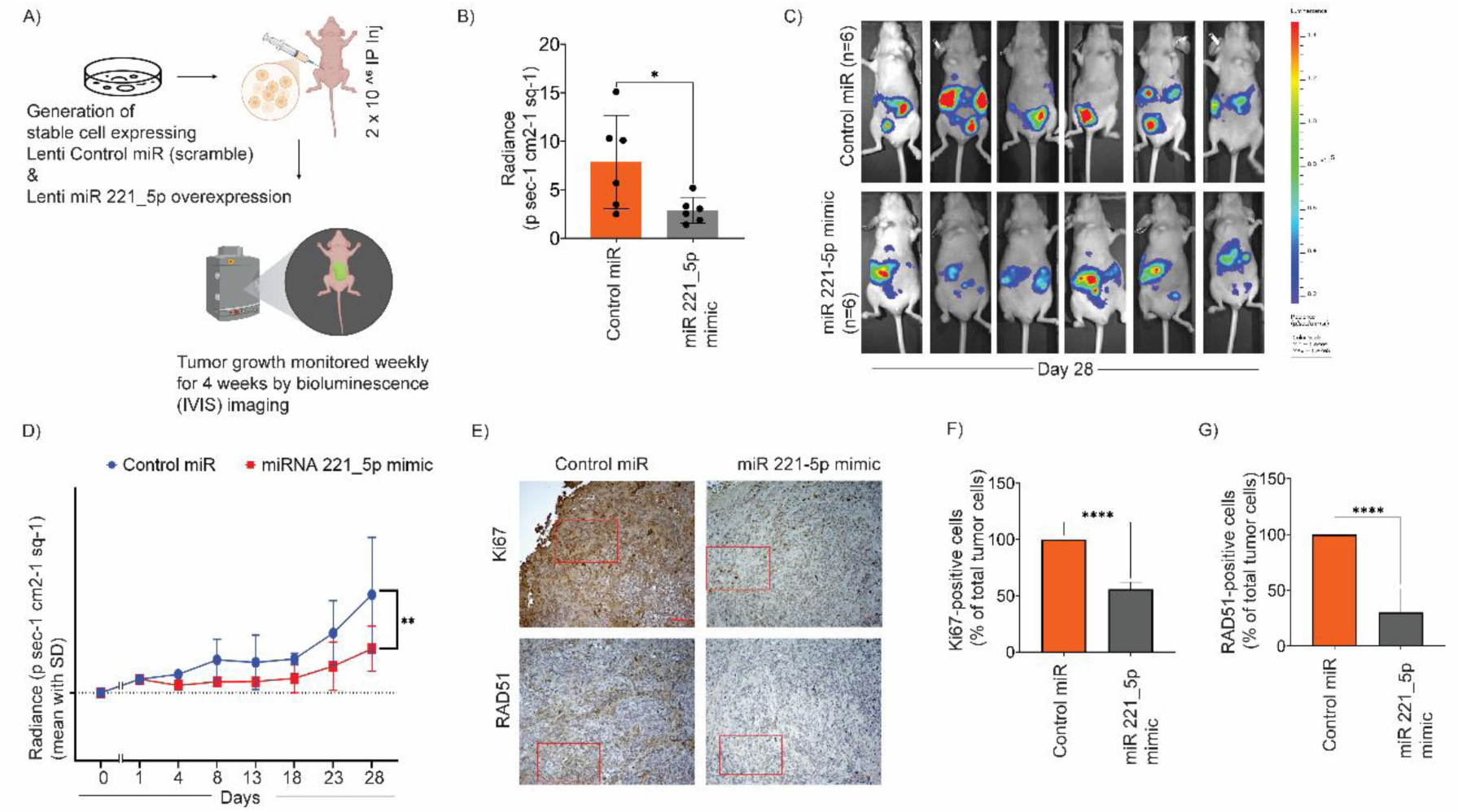
Stable miR–221–5p expression attenuates ovarian tumor growth and knocks down RAD51 in vivo. (A) Schematic of the experimental workflow. Athymic nude mice (n = 6/group) were intraperitoneally (i.p.) injected with SKOV3-Luc cells stably expressing either a control miRNA or miR-221-5p. Tumor progression was monitored weekly starting at day 10 post-implantation using bioluminescence imaging (IVIS). (B) Quantification of total radiance (photons s⁻¹ cm⁻² sr⁻¹) measured on day 28 endpoint.Data represents mean ± SD; n = 6 mice per group; *P < 0.05, two–tailed *t*–test). (C) Representative IVIS bioluminescence images of mice from the control and miR-221-5p groups on day 28. The pseudocolor scale indicates the radiance intensity, reflecting tumor burden. (D) Longitudinal tumor growth curves based on bioluminescence radiance over time. Statistical significance was determined by two-way repeated-measures ANOVA (mean ± SD; **P < 0.01). (E) Representative immunohistochemistry (IHC) images of harvested tumor sections stained for the proliferation marker Ki67 (top) and the DNA repair protein RAD51 (bottom). Scale bars, 100 μm. (F, G) Quantitative analysis of Ki67–positive (F) and RAD51–positive (G) cells, expressed as a percentage of total cells per high-power field (n = 3 tumors per group; data represented mean ± SD; (****P < 0.0001, two–tailed *t*–test).

Following molecular characterization, cells were implanted intraperitoneally (IP) into nude mice (n=6; per group) to model the clinical dissemination characteristic of advanced-stage ovarian cancer. Tumor progression was monitored non-invasively using bioluminescence imaging (IVIS). To allow for stable engraftment, longitudinal bioluminescence measurements were initiated 10 days post-implantation (Fig. 6A). Quantitative analysis of the bioluminescent signal intensity revealed a significant and progressive decrease in tumor burden in mice harboring miR-221-5p-expressing cells compared to the control cohort (Fig. 6B–6D). Statistical validation using Two-way ANOVA, incorporating both time and tumor growth variables, confirmed a highly significant reduction in the overall tumor burden in the miR-221-5p group.

To correlate these physiological outcomes with our proposed molecular mechanism, harvested tumor tissues were analyzed via immunohistochemistry (IHC). Consistent with our *in vitro* findings, miR-221-5p overexpression resulted in a marked depletion of RAD51 protein levels within the tumor parenchyma (Fig. 6E–6G). Furthermore, staining for the proliferation marker Ki67 revealed significantly diminished proliferative activity in miR-221-5p-expressing xenografts compared to controls (Fig. 6E–6F). Collectively, these results demonstrate that the restoration of miR-221-5p effectively suppresses ovarian tumor growth and dissemination *in vivo* by collapsing the DNA repair axis and limiting cellular proliferation.

### miR-221-5p Restoration Overcomes Acquired Platinum Resistance and Induces a Synthetic Lethal Vulnerability to PARP Inhibition

To determine whether the RAD18/RAD51 axis actively drives the development of platinum resistance, we utilized isogenic pairs of platinum-sensitive (A2780) and acquired-resistant (A2780/CP70) ovarian cancer cell lines. Western blot analysis revealed substantially elevated levels of both RAD18 and RAD51 in the resistant A2780/CP70 cells compared to their sensitive parental counterparts and A2780 cells (Fig. 7A). This upregulation of the repair axis corresponded to a significantly higher clonogenic survival capacity in A2780/CP70 cells across all tested carboplatin doses (Fig. 7B–7C). Consistent with our proposed model, ectopic restoration of miR-221-5p in these resistant lines effectively depleted RAD18 and RAD51 protein levels (Fig. 7D). Notably, miR-221-5p expression significantly re-sensitized the resistant A2780/CP70 cells to carboplatin, resulting in a dramatic reduction in colony-forming ability that effectively mirrored the sensitivity of the parental sensitive line (Fig. 7E–F).

**Figure 7.**
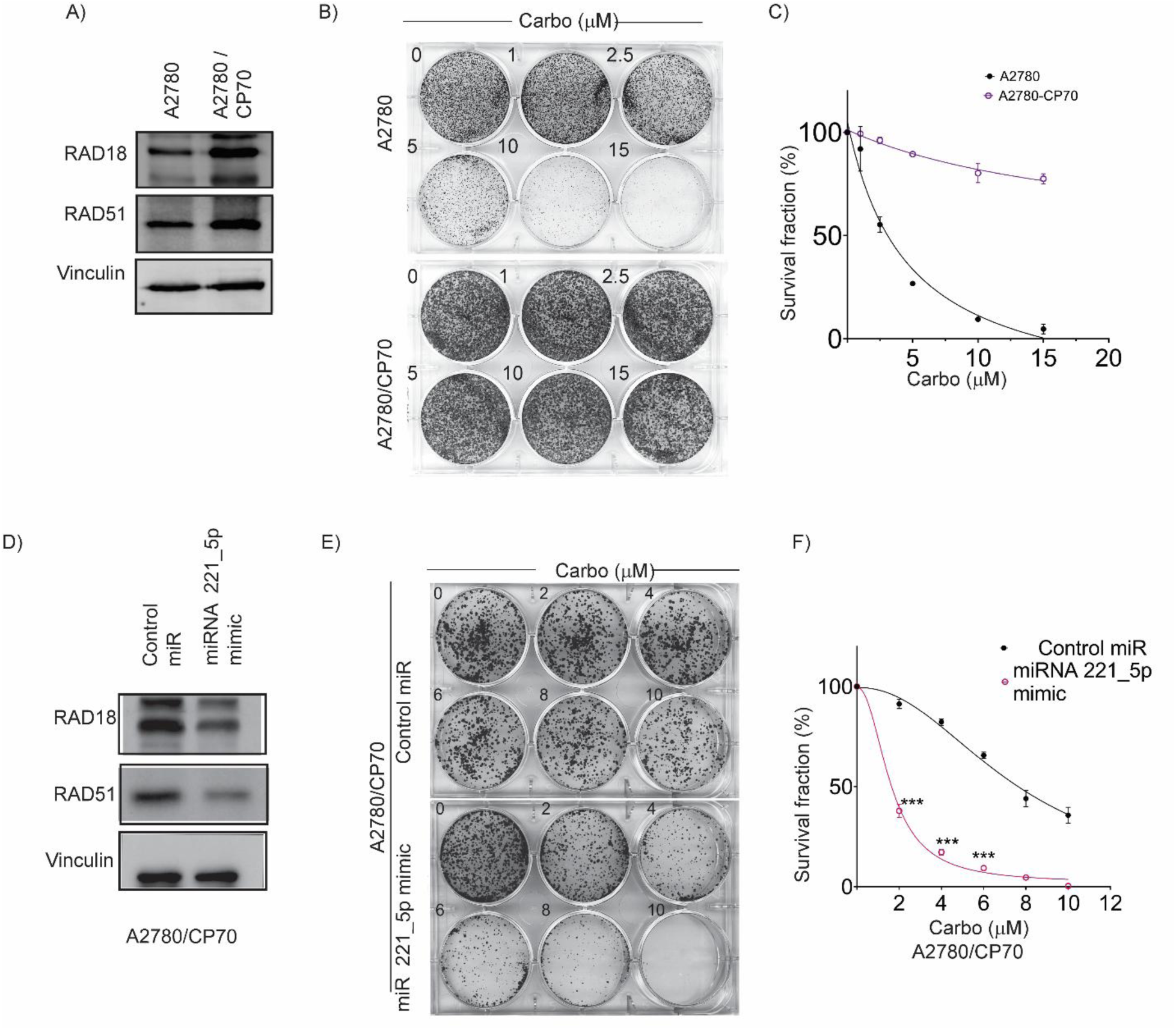
RAD18/RAD51 upregulation confers platinum resistance, and miR-221-5p restoration resensitizes resistant cells. (A) Immuno blot analysis of RAD18 and RAD51 expression in platinum-sensitive A2780 cells and their platinum-resistant isogenic counterpart A2780/CP70. Vinculin used as a loading control. (B) Representative images (B) and corresponding survival fraction curves (C) from clonogenic survival assays of A2780 and A2780/CP70 cells treated with increasing concentrations of carboplatin (0–15 μM). Data illustrates the significant shift in the between sensitive and resistant lines (mean ± SD, n = 3). (D) Immuno blot showing the depletion of RAD18 and RAD51 in platinum-resistant A2780/CP70 cells 48 h post-transfected with a control miR or a miR-221-5p mimics. (E, F) Representative images (E) and quantitative survival fraction analysis (F) of a low-density colony formation assay in A2780/CP70 cells. Restoration of miR-221-5p significantly sensitized resistant cells to a carboplatin gradient (0–10 μM). Data are presented as mean ± SD from three independent biological replicates (***P < 0.001, two–tailed t–test). (***P < 0.001, two–tailed t–test).

Since RAD18 is essential for recruiting RAD51 to stalled or collapsed replication forks to facilitate HR repair[33][34], we hypothesized that miR-221-5p-mediated depletion of these factors would induce a functional HR-deficiency (HRD) or “BRCAness-like” phenotype. This state typically confers an increased vulnerability to PARP inhibitors (PARPi) via synthetic lethality. In OVCAR8, A2780 and SKOV3 cells, treatment with the PARPi Olaparib significantly reduced clonogenic survival in a dose-dependent manner specifically when combined with miR-221-5p mimics (Fig. 8A–C and supplemental figure 8A–8B).

**Figure 8.**
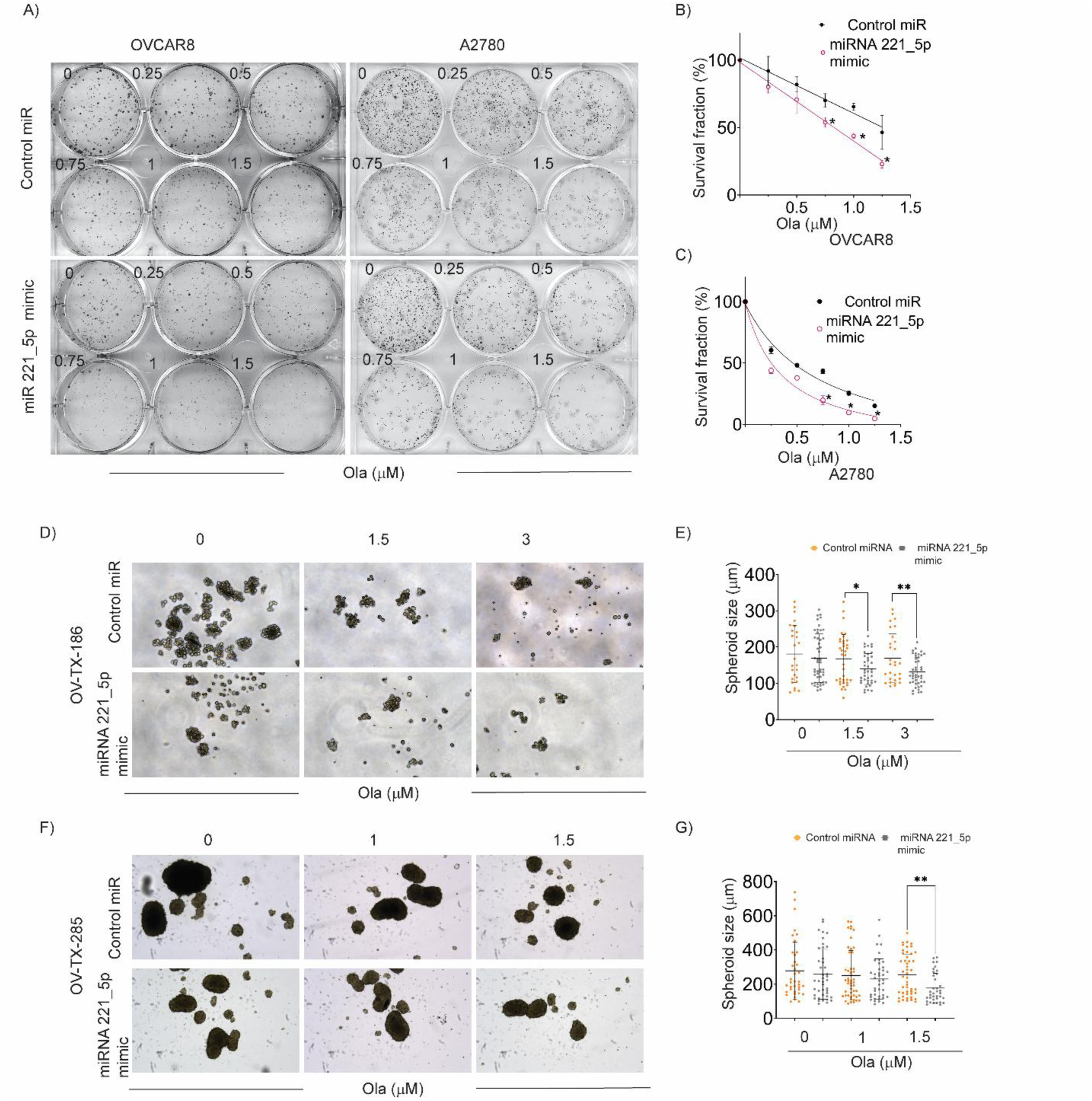
miR-221-5p restoration induces synthetic lethality with Olaparib in OC cell lines and patient-derived organoids. (A) Representative images of low-density clonogenic assays. OVCAR8 (left) and A2780 (right) cells were transfected with control miRNA or miR-221-5p mimics and treated with increasing concentrations of the PARP inhibitor Olaparib (0–1.5 μM). (B,C) Survival dose–response curves quantifying colony-forming efficiency (normalized to untreated controls) for OVCAR8 (B) and A2780 (C). Data are mean ± SD of three independent experiments. (*P < 0.05, **P < 0.01; two–tailed *t*–test). (D,E) Evaluation of the miR-221-5p/Olaparib interaction in the Patient-derived organoid model OV-TX-186. Representative brightfield images (D) and corresponding violin plots (E) showing organoids size distribution following miR-221-5p mimic transfection and olaparib treatment (0, 1.5, 3 μM). (F,G) Validation in the patient-derived organoid model OV-TX-285. Representative images (F) and quantitative violin plots (G) displaying organoid size under the indicated conditions in (D,E). For organoid analysis, approximately 50 organoids per condition were measured. Organoids with a 50 μm diameter were included in the quantification. Data are presented as mean ± SD of three biological replicates Organoid size greater than 50 μm were counted (**P < 0.01, ***P < 0.001; two–tailed *t*–test;).

To confirm this therapeutic synergy in a more physiologically relevant system, we tested the combination in primary patient-derived organoids (OV-TX-186 and OV-TX-285). Consistent with our cell line data, miR-221-5p restoration markedly sensitized primary organoids to Olaparib, resulting in a significant decrease in organoid growth and survival compared to control-treated primary cells (Fig. 8D–G).

Collectively, these results demonstrate that the upregulation of RAD18 and RAD51 is a hallmark of acquired platinum resistance in OC. By repressing this dual repair axis, miR-221-5p restoration not only overcomes platinum resistance but also creates a new therapeutic window for PARP inhibition, highlighting its potential as a precision oncology strategy for aggressive, HR-proficient ovarian cancer.

## Discussion

Ovarian cancer (OC) remains the most lethal gynecological malignancy[35][36], primarily due to the rapid emergence of acquired platinum resistance[37][34]. While initial surgical debulking and platinum-based chemotherapy are often clinically successful, the inherent plasticity of the DNA damage response (DDR) allows tumor cells to rewire survival pathways, inevitably leading to relapses. In this study, we identify the miR-221-5p/RAD18/RAD51 axis as a critical molecular determinant of this resistance[38][39][40]. Our findings demonstrate that the loss of miR-221-5p acts as an epigenetic “release” that permits the simultaneous upregulation of two distinct yet synergistic repair pathways: translesion synthesis (TLS) via RAD18 and homologous recombination (HR) via RAD51.

The prognostic significance of RAD51 and RAD18 in OC is well-supported by our TCGA analysis and established clinical literature[38][41][42]. RAD51 is the central recombinase of HR; recent studies have utilized quantitative immunohistochemistry to demonstrate that high RAD51 expression is a “bona fide” biomarker for early relapse and poor overall survival in epithelial ovarian cancer (EOC) [43]. Furthermore, RAD51 overexpression is a frequent feature of high-grade serous ovarian cancer (HGSOC) and is increasingly associated with the reversal of “functional BRCAness”, where tumors restore HR competency to evade platinum-induced lethality even in the absence of *BRCA1/2* reversion mutations[38][44].

Similarly, RAD18 has emerged as a cornerstone of DNA damage tolerance. By orchestrating the mono-ubiquitination of PCNA, RAD18 enables replication forks to bypass bulky platinum-DNA adducts via TLS, preventing lethal fork collapse and subsequent apoptosis [3]. Our observation that both proteins are significantly overexpressed in platinum-resistant isogenic pairs (A2780/CP70 and SKOV3/CP20) reinforces the concept that resistant OC cells do not rely on a single repair mechanism. Instead, they employ a coordinated “dual-pathway” network that integrates TLS and HR to survive the intense genotoxic stress imposed by standard-of-care therapies[2][45][46].

A central novelty of this work is the characterization of miR-221-5p as a master regulator of this coordinated repair response. While the miR-221/222 cluster is traditionally categorized as oncogenic and largely based on studies of the miR-221-3p arm and its suppression of the p27Kip1 tumor suppressor [47][48]. Our data clearly defines the -5p strand as a potent tumor suppressor in OC. This “arm switching” or functional divergence between miRNA strands is an emerging and complex theme in RNA biology, where the selection of the mature strand (5p vs. 3p) can dictate entirely opposite cellular fates [49][17]. We provide definitive evidence that miR-221-5p directly binds the 3′UTRs of both *RAD18* and *RAD51*, leading to their post-transcriptional silencing. This direct regulation was functionally validated by the reduction of critical downstream signaling markers, specifically Ub-PCNA and Ub-FANCD2, indicating a systemic failure of the cell’s damage tolerance machinery.

The functional consequences of miR-221-5p restoration are striking. By collapsing the RAD18/RAD51 axis, we effectively induced a state of homologous recombination deficiency (HRD), as evidenced by diminished DR-GFP reporter activity and attenuated RAD51 foci formation. This molecular collapse was accompanied by the accumulation of persistent 53BP1 foci and increased genomic instability, as shown by alkaline comet assays. Crucially, our flow cytometry data revealed that these effects were independent of cell cycle redistribution. It is imperative to rule out the contribution of any cell cycle discrepancies in miR-221-5P-transfected cells to RAD51 and RAD18 downregulation as non-replicating cells tend to have low expression of the HR proteins. This suggests that miR-221-5p does not merely act as a cytostatic agent but functions as a direct chemosensitizer that lowers the threshold for DNA-damage-induced cell death.

From a translational perspective, the ability of miR-221-5p to resensitize resistant cells in 3D patient-derived organoids and *in vivo* xenografts highlights its therapeutic potential. Currently, PARP inhibitors (PARPi) are the standard of care for HRD-positive tumors; however, approximately 50% of HGSOC patients are HR-proficient and show disappointing responses to PARPi [19]. Our data suggest that miR-221-5p replacement therapy could serve as a “priming” strategy to induce a transient HRD phenotype in otherwise proficient tumors. This “synthetic lethal” approach would potentially expand the cohort of patients eligible for PARPi or enhance the efficacy of platinum-based agents in the resistant setting.

## Conclusion

Our study delineates a previously unrecognized epigenetic regulatory axis in ovarian cancer. We propose that the downregulation of miR-221-5p is a primary driver of acquired chemoresistance through the derepression of RAD18 and RAD51. Restoring miR-221-5p disrupts the coordinated TLS and HR repair pathways, promoting genomic instability and overcoming the formidable barrier of platinum resistance. These findings provide a mechanistic rationale for the development of miR-221-5p mimics as a precision medicine intervention to combat aggressive, chemoresistant ovarian cancer.

## Materials and Methods

### Cell lines, Culture Conditions and Reagents

The human epithelial ovarian cancer (OC) cell lines OVCAR3, OVCAR4, OVCAR8, OV90, ES2, A2780, and SKOV3 were purchased from the American Type Culture Collection (ATCC; Manassas, VA). To model acquired chemoresistance, isogenic platinum-sensitive/resistant pairs (A2780 vs. A2780/CP70 and SKOV3 vs. SKOV3/CP20) were utilized as previously described [50]. Immortalized human fallopian tube epithelial cells (FTEC) served as a non-malignant control [51].

All cell lines were cultured in their respective vendor-recommended media (e.g., DMEM, RPMI-1640, or DMEM/F12) supplemented with 10% fetal bovine serum (FBS; Omega Scientific Inc.) and 1% penicillin-streptomycin (Invitrogen). Cultures were maintained in a humidified atmosphere at 37°C incubator with 5% CO2. All lines were routinely screened for *Mycoplasma* contamination using PCR-based detection (Cat No: AB289834). Carboplatin (Carbo; Selleckchem) was dissolved in ultrapure water to create stock solutions and applied at the concentrations and durations specified in the figure legends.

### Bioinformatics and Clinical Correlation Analysis

#### Clinical Survival Analysis

The prognostic value of *RAD18* and *RAD51* mRNA expression was evaluated using the Kaplan-Meier (KM) Plotter database [27]. We analyzed a combined cohort of 614 ovarian cancer patients derived from The Cancer Genome Atlas (TCGA) and GSE14764 datasets. Patients were stratified into high- or low-expression cohorts using the median transcript level as the discriminatory cutoff. Overall survival (OS) and progression-free survival (PFS) were compared using the log-rank test. Survival outcomes are reported as hazard ratios (HR) with 95% confidence intervals (CI) and corresponding *p* values.

#### Differential miRNA Expression Analysis

To identify clinically relevant miRNAs, we re-analyzed the Gene Expression Omnibus (GEO) dataset GSE235525[28], comparing miRNA expression profiles from the serum of patients with high-grade serous ovarian cancer (HGSOC) against healthy controls. Differentially expressed miRNAs (DEMs) were identified using the GEO2R interactive tool. Statistical significance was defined by an adjusted *P* < 0.05 and a log2 fold-change > 1.

#### *In silico* Gene Expression Validation

The differential expression of *RAD18* and *RAD51* between malignant ovarian tissues and normal controls was validated using the GEPIA (Gene Expression Profiling Interactive Analysis) portal[26]. This analysis integrated transcriptomic data from TCGA (tumor) and the Genotype-Tissue Expression (GTEx) project (normal) using a standardized processing pipeline.

### Dual-Luciferase 3’UTR Reporter Assay

To confirm the direct post-transcriptional regulation of *RAD18* and *RAD51* by miR-221-5p, individual GoClone 3′ UTR reporter constructs (Active Motif, Carlsbad, CA) were utilized. These vectors contain the firefly luciferase open reading frame (ORF) fused to the 3′ UTR regions of *RAD18* or *RAD51* harboring the predicted miR-221-5p binding sites.

ES2 and OVCAR8 cells were seeded in 96-well plates and co-transfected with 100 ng of the respective luciferase reporter vectors and 50 nM of either miR-221-5p mimics (Cat# C-301163-01-0020) or non-targeting control mimics (Cat# CN-001000-01-20; Horizon Discovery) using Lipofectamine RNAiMAX (Life Technologies). After 48 h of incubation, luciferase activity was quantified using the LightSwitch Assay System (SwitchGear Genomics) on a Tecan microplate reader according to the manufacturer’s instructions. To account for variations in transfection efficiency, firefly luciferase activity was normalized to either a co-transfected constitutive reporter or to total protein concentration (as per the LightSwitch system protocol). Results are expressed as relative luciferase activity compared to the non-targeting control.

### DNA Repair and Functional Assays

#### Homologous Recombination (HR) Reporter Assay

To quantify HR efficiency, a functional DR-GFP reporter system was utilized as previously described [52]. The reporter consists of two non-functional GFP derivatives: a SceGFP gene containing an I-SceI recognition site and an internal stop codon, and a truncated iGFP fragment. The pDR-GFP and pCBASceI plasmids were kindly provided by Dr. Maria Jasin (Addgene plasmids #26477 and #26475).

OVCAR8 cells stably harboring the pDR-GFP cassette were transfected with the pCBASceI expression vector to induce a site-specific DNA double-strand break (DSB). Cells were simultaneously transfected with miR-control or miR-221-5p mimics. Successful HR-mediated repair of the DSB using the iGFP fragment as a template restores the functional GFP open reading frame. At 48 h post-transfection, the frequency of GFP-positive cells was quantified via flow cytometry (BD Accuri) to determine the relative HR efficiency.

#### Alkaline Comet Assay

To assess global DNA damage and the persistence of strand breaks, an alkaline comet assay (single-cell gel electrophoresis) was performed using the OxiSelect 96-well Comet Assay Kit (Cell Biolabs, San Diego, CA). Briefly, cells were treated with 5 μM carboplatin for 24 h, harvested, and embedded in 1% low-melting-point agarose on specialized glass slides.

To permit the detection of both single- and double-strand breaks, the cells were subjected to alkaline lysis followed by DNA unwinding in an alkaline electrophoresis buffer (pH > 13). Electrophoresis was conducted at 1 V/cm for 30 min to allow damaged DNA to migrate from the nucleus, forming a “comet tail.” Following neutralization and ethanol dehydration, DNA was stained with Vista Green DNA Dye. Comets were visualized using a Zeiss Axio fluorescence microscope, and genomic damage was quantified by measuring the tail moment and tail DNA percentage in at least 50 randomly selected cells per condition using ImageJ software (NIH) equipped with the Open Comet plugin.

DNA damage was assessed under alkaline conditions using the OxiSelect 96-well Comet Assay Kit (Cell Biolabs). Cells were treated with 5 μM carboplatin for 24 h, immobilized in 1% low-melting-point agarose, and subjected to electrophoresis. After neutralization and staining with Vista Green DNA dye, comets were imaged (Zeiss Axio) and analyzed using the ImageJ comet plugin.

#### Clonogenic Survival Assay

Cellular viability and proliferative capacity following treatment were assessed via clonogenic assay. Cells were seeded at low density (500–5,000 cells/well) in 6-well plates and allowed to adhere overnight. Cells were subsequently treated with indicated concentrations of carboplatin or olaparib, and cells were cultured for 7–10 days to allow for colony formation. Once control wells reached optimal density, colonies were fixed with methanol and stained with 1% crystal violet solution.

Colonies consisting of ≥ 50 cells were manually or semi-automatically quantified using ImageJ software (NIH, Bethesda, MD). The surviving fraction was calculated by normalizing the plating efficiency of treated groups to that of the respective non-targeting control (miR-NC) or vehicle-treated groups.

### 3D Culture and Invasion Assays

#### Patient-derived 3D Organoid Cultures

Primary ovarian cancer (OC) cells (OV-TX-186 and OV-TX-285), obtained from the Texas Tech University Health Sciences Center (TTUHSC) Cancer Center Biorepository, were utilized to generate patient-derived organoids. To promote anchorage-independent growth, cells were seeded in ultra-low attachment (ULA) 6-well plates (Corning). The organoids were maintained in DMEM/F12 supplemented with 20% FBS, 1X B-27, 1X N-2, 20 ng/mL EGF, and 20 ng/mL FGF. Organoid morphology and assembly were monitored for 7–14 days. Quantitative analysis of organoid diameter and volume was performed using Nikon NIS-Elements (ND2) software.

#### 3D Matrix-Dependent Invasion Assay

To evaluate metastatic potential, multicellular spheroids were first generated in low-adhesion round-bottom 96-well plates at a density of 3,000 cells/well. Following a 72-h incubation to allow compact spheroid formation, 50 L of Cultrex 3D Spheroid Invasion Matrix (R&D Systems) was overlaid. The invasive capacity of the spheroids was monitored over 3–6 days using a Nikon Eclipse TE confocal microscope. The degree of invasion was determined by measuring the invasive area and perimeter expansion relative to the day 0 core using ImageJ software (NIH).

### Molecular Analysis

#### Western Blotting Analysis

Total cellular proteins were extracted using RIPA lysis buffer (Thermo Fisher Scientific) supplemented with a protease and phosphatase inhibitor cocktail (Roche, Basel, Switzerland). Protein concentrations were determined using the Bradford assay (Bio-Rad). Equal amounts of protein were denatured and resolved by SDS-PAGE using 8–12% gradient gels.

The resolved proteins were then electro-transferred onto nitrocellulose membranes (GE Healthcare). Membranes were blocked for 1 h at room temperature with 5% non-fat dry milk in Tris-buffered saline containing 0.1% Tween-20 (TBST) to minimize non-specific binding. Membranes were subsequently incubated overnight at 4°C with indicated primary antibodies.

Following three washes with TBST, membranes were incubated with horseradish peroxidase (HRP)-conjugated secondary antibodies for 1 h at room temperature. Protein bands were visualized using enhanced chemiluminescence (ECL) reagents (PerkinElmer) and captured on an Azure C300 imaging system (Azure Biosystems) or using X-ray film developer. Densitometric quantification was performed using **ImageJ software**, with target protein levels normalized to **vinculin**, β**-actin**, or **GAPDH** as internal loading controls.

#### Quantitative Real Time PCR (qRT-PCR

Total RNA was isolated using the Quick-RNA MiniPrep kit (Zymo Research, Irvine, CA) according to the manufacturer’s instructions. RNA concentration and purity were assessed using a NanoDrop spectrophotometer. For mRNA quantification, cDNA was synthesized from of total RNA using the High-Capacity cDNA Reverse Transcription Kit (Invitrogen) with random hexamer primers. For miRNA quantification, cDNA was generated using the Mir-X miRNA First-Strand Synthesis Kit (TaKaRa Bio, Mountain View, CA), which utilizes a poly-A tailing strategy.

Quantitative PCR was performed using PowerUp SYBR Green Master Mix (Applied Biosystems) on a QuantStudio 12K Flex Real-Time PCR System. Specificity of the amplification was confirmed by melt-curve analysis. Relative gene expression was calculated using the 2−ΔΔCt method. Target mRNA levels were normalized to *GAPDH*, while miRNA levels were normalized to the small nuclear RNA *U6*.

#### Cell Cycle Distribution Analysis

Cell cycle progression was evaluated via flow cytometry using propidium iodide (PI) staining. Following transfection with miR-control or miR-221-5p mimics, cells were harvested by trypsinization, washed twice with ice-cold PBS, and fixed in 100% ice-cold ethanol at -20^0^ C overnight.

Fixed cells were subsequently washed with PBS to remove residual ethanol and resuspended in a staining solution containing 50 µg/mL PI (Invitrogen) and 100 µg/mL RNase A (Thermo Scientific) to ensure DNA-specific binding. After a 30-minute incubation in the dark at room temperature, DNA content was analyzed using a BD Accuri C6 flow cytometer (BD Biosciences). A minimum of 10,000 events were captured per sample. The percentages of cells in G1, S and G2/M phases were quantified using ModFit LT 5.0 software (Verity Software House).

### Immunofluorescence and Confocal Microscopy

#### 2D Monolayer Imaging

Cells were seeded into FluoroDishes (World Precision Instruments) and allowed to adhere overnight. Following treatment, cells were fixed with 4% paraformaldehyde for 10 min at room temperature and permeabilized with 0.2% Triton X-100 in PBS for 3 min. To minimize non-specific binding, cells were blocked with 10% normal goat serum for 40 min. Samples were incubated overnight at 4°C with primary antibodies against 53BP1 (1:500; Cell Signaling, Cat# 4937) or RAD51 (1:200; Millipore, Cat# 65653). Following PBS washes, cells were incubated with Alexa Fluor-conjugated secondary antibodies (1:500; Molecular Probes) for 2 h at room temperature. Nuclei were counterstained and mounted using Vectashield with DAPI (Vector Laboratories).

#### 3D Spheroid and Organoid Imaging

For 3D cultures, spheroids were fixed in 4% PFA for 1 h, followed by extended permeabilization with 0.2% Triton X-100 for 20 min. After blocking, spheres were incubated overnight with anti-RAD18 (1:200; Cell Signaling, Cat# 9040) or RAD51 (1:200; Millipore, Cat# 65653). To visualize the 3D architecture and cytoskeleton, spheres were stained with fluorescent phalloidin (Invitrogen, Cat# A12379) for 1 h.

All images were captured using a Nikon Eclipse TE confocal microscope equipped with a high-resolution Z-axis motor. For 3D analysis, Z-stacks were acquired at 5.0 µm intervals and processed as maximum intensity projections. Foci formation (defined as ≥ 5 foci per nucleus) and integrated fluorescence intensity were quantified using ImageJ software (NIH).

### Animal Studies and *In Vivo* Xenograft Model

#### Lentiviral Vector Transduction and Stable Cell Line Generation

To generate stable miR-221-5p expression, SKOV3-Luc cells (expressing firefly luciferase) were seeded at a density of 5 X 10^4^ cells/cm^2^. After 24 h, cells were transduced with lentiviral vectors encoding either miR-221-5p (pLenti-III-miR-GFP) or a non-targeting control (pLenti-III-miR-GFP-Blank; Applied Biological Materials, Richmond, BC, Canada) at a multiplicity of infection (MOI) of 15. Viral supernatant was replaced with complete growth medium 24 h post-transduction. Transduction efficiency was validated by monitoring GFP expression via fluorescence microscopy and confirmed by Western blot analysis of target repair proteins prior to implantation.

#### Intraperitoneal (i.p.) Xenograft Model

All animal experiments were performed in accordance with the NIH Guide for the Care and Use of Laboratory Animals and ARRIVE guidelines, under protocols approved by the TTUHSC Institutional Animal Care and Use Committee (IACUC). Female athymic nude mice (8 weeks old) were randomized into two groups (n = 6 per group). Mice received an intraperitoneal (i.p.) injection of 2X 10^6^ SKOV3-Luc cells (control vs. miR-221-5p). Tumor progression and peritoneal dissemination were monitored 2–3 times weekly using bioluminescence imaging (IVIS; PerkinElmer). Mice were injected i.p. with D-luciferin (150 mg/kg) and imaged under isoflurane anesthesia, and bioluminescence intensity was quantified as total radiance (photons/s/cm^2^/sq). At the experimental endpoint (28–32 days post-injection), mice were euthanized, and tumors were harvested, weighed, and fixed in 4% paraformaldehyde for subsequent immunohistochemistry (IHC) analysis.

## Statistical Analysis

Statistical analyses were performed using GraphPad Prism v.8.4.3 (GraphPad Software, San Diego, CA). Quantitative data are presented as the mean ± standard deviation (SD) from at least three independent biological replicates, unless otherwise specified. For comparisons between two experimental groups, a two-tailed unpaired Student’s *t*-test was employed, assuming equal variance. For experiments involving multiple groups or treatment conditions, one-way or two-way analysis of variance (ANOVA) followed by Tukey’s or Sidak’s post-hoc test for multiple comparisons was used. Longitudinal tumor growth data from IVIS imaging were analyzed using two-way repeated-measures ANOVA. A *P*-value < 0.05 was considered statistically significant. Significance levels are denoted in the figures as follows: ∗P<0.05, ∗∗P<0.01, ∗∗∗P<0.001, ∗∗∗∗P<0.0001).

## Author Contributions

Conceived and designed the experiments: TO, MBR, CM and KP. Performed the experiments: TO, NS, SK and AI. Wrote the paper: TO and KP. Reviewed and edited the manuscript: All the authors analyzed the data, reviewed and edited the manuscript.

## Acknowledgments

We thank all the lab members of KP for their thoughtful discussion and suggestions. This work is supported by grants from NIH CA219187 (KP), Weitlauf Endowment for Cancer Research (KP), Rural Cancer Collaborative Program seed grant (KP), and Laura W. Bush Foundation for Women’s Health (MBR and KP). We thank Dr. Reynolds for providing patient-derived ovarian cancer models used in this study.

