## Supplemental figures for "The miR-221-5p/RAD18/RAD51 Axis Regulates DNA Damage Tolerance and Homologous Recombination to Drive Platinum Resistance in Ovarian Cancer"

### SUPPLEMENATRY FIGURE LEGENDS

**Supplementary Figure 1. miR-221-5p is markedly downregulated in ovarian cancer patients and cell lines, and its restoration suppresses OC cell clonogenic potential and spheroid growth.**

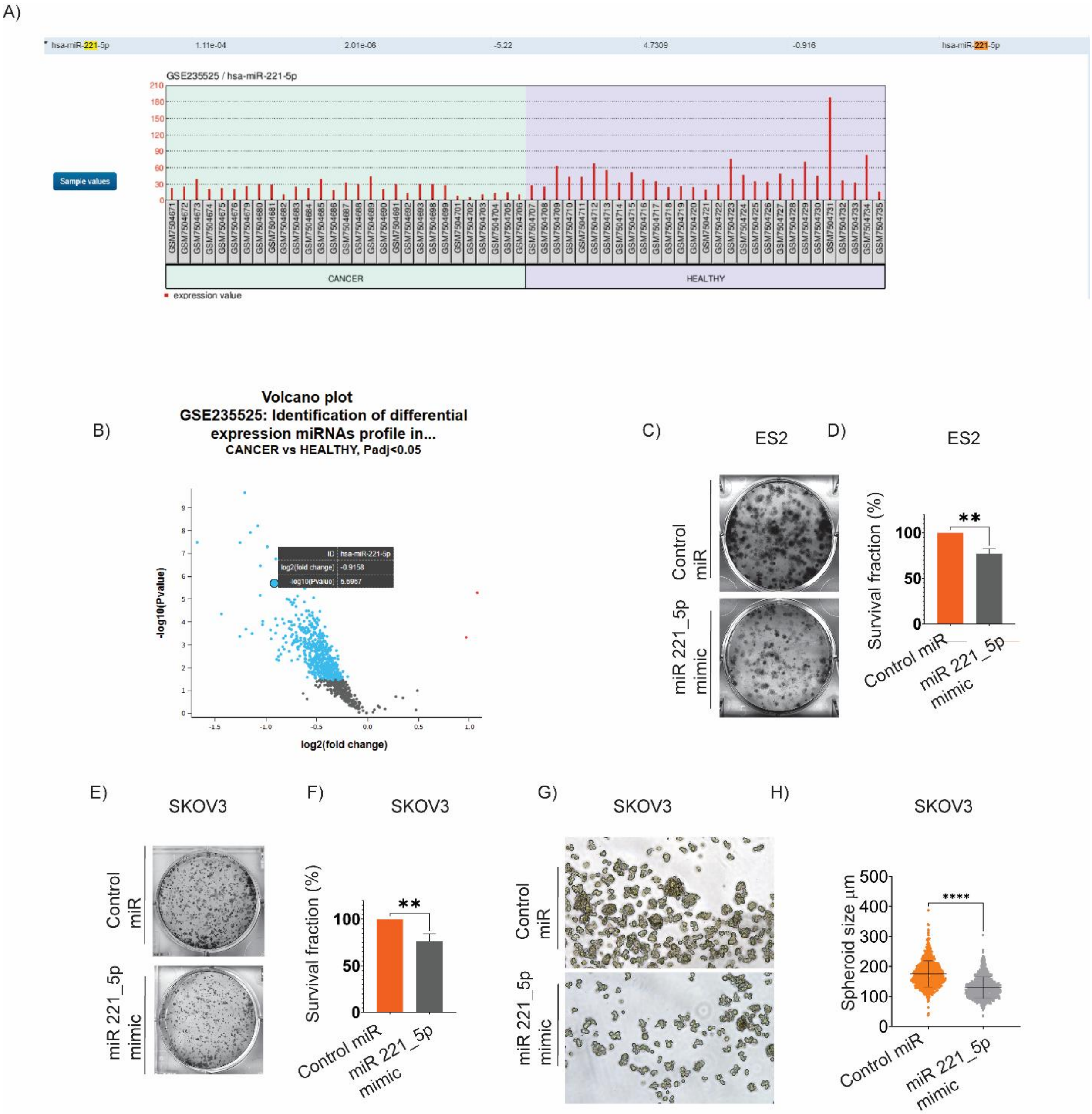


(A) Re-analysis of the GEO dataset GSE235525 comparing serum miRNA expression profiles from HGSOC patients (CANCER) versus healthy controls (HEALTHY). Bars represent individual sample expression values of *hsa-miR-221-5p* (red) ($\log_2$ fold change ≈ −5.22; adjusted P = 1.1 × $10^{-4}$).

Volcano plot illustrates differentially expressed miRNAs in the GSE235525 cohort (cancer vs. healthy). Each point denotes a miRNA; blue points indicate significantly downregulated miRNAs, while red points indicate significantly upregulated miRNAs. The position of miRNA-221-5p is indicated (P adj < 0.05).

(B,C) Representative images (B) and quantitative survival fraction analysis (C) of low-density colony formation assays in ES2 cells 10 days post-transfection with control miRNA or miR-221-5p mimics. Data presented is mean ± SD of three independent experiments (**P < 0.01; mean ± SD). (**P < 0.01; mean ± SD).

(D,E) Representative images (E) and quantitative survival fraction analysis (F) of clonogenic assays in SKOV3 cells following miR-221-5p restoration (**P < 0.01; n = 3).

(F,G) Representative images (F) and quantitative violin plots (G) of 3D spheroid cultures of SKOV3 cells transfected with Control miR or miR-221-5P mimic. Data presented represents mean ± SD of three independent experiments. Spheroids bigger than 50 µm were calculated for plotting. (****P < 0.0001, two-tailed *t*-test).

**Supplementary Figure 2. miR-221-5p mimic depletes RAD18 and RAD51 transcripts and validates 3′-UTR reporter mechanism.**

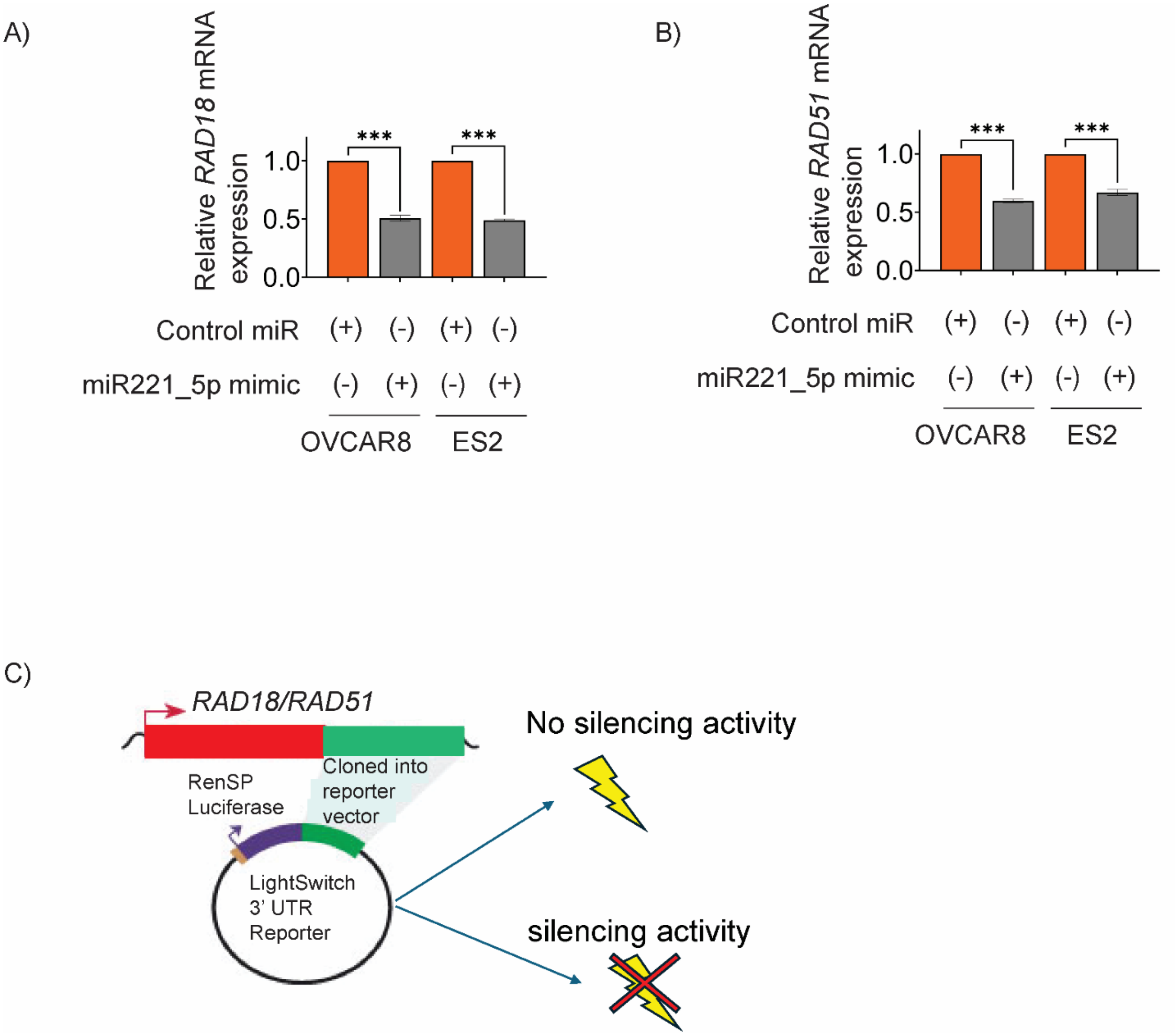


(A,B) Quantitative RT-PCR analysis of *RAD18* (A) and *RAD51* (B) mRNA levels in OVCAR8 and ES2 cells 24 h post-transfection with Control miR or miR-221-5P mimics. Transcript levels were normalized to *GAPDH.* Data represent mean ± SD of three independent experiments. (***P < 0.001, two-tailed *t*-test).

(C) Schematic representation of the LightSwitch 3′-UTR luciferase reporter assay strategy used to validate direct targeting engagement. The wildtype 3′-UTR of *RAD18* and *RAD51* was cloned downstream of the RenSP luciferase reporter gene. In the absence of miR-221-5p (Control), high luciferase activity is maintained ("no silencing activity"). Upon miR-221-5p binding to its complementary seed sites within 3'UTR, luciferase expression is suppressed ("silencing activity"). This mechanistic framework serves as the basis for the dual-luciferase results shown in Figure 2.

**Supplementary Figure 4. miR-221-5p restoration does not alter cell-cycle distribution.**

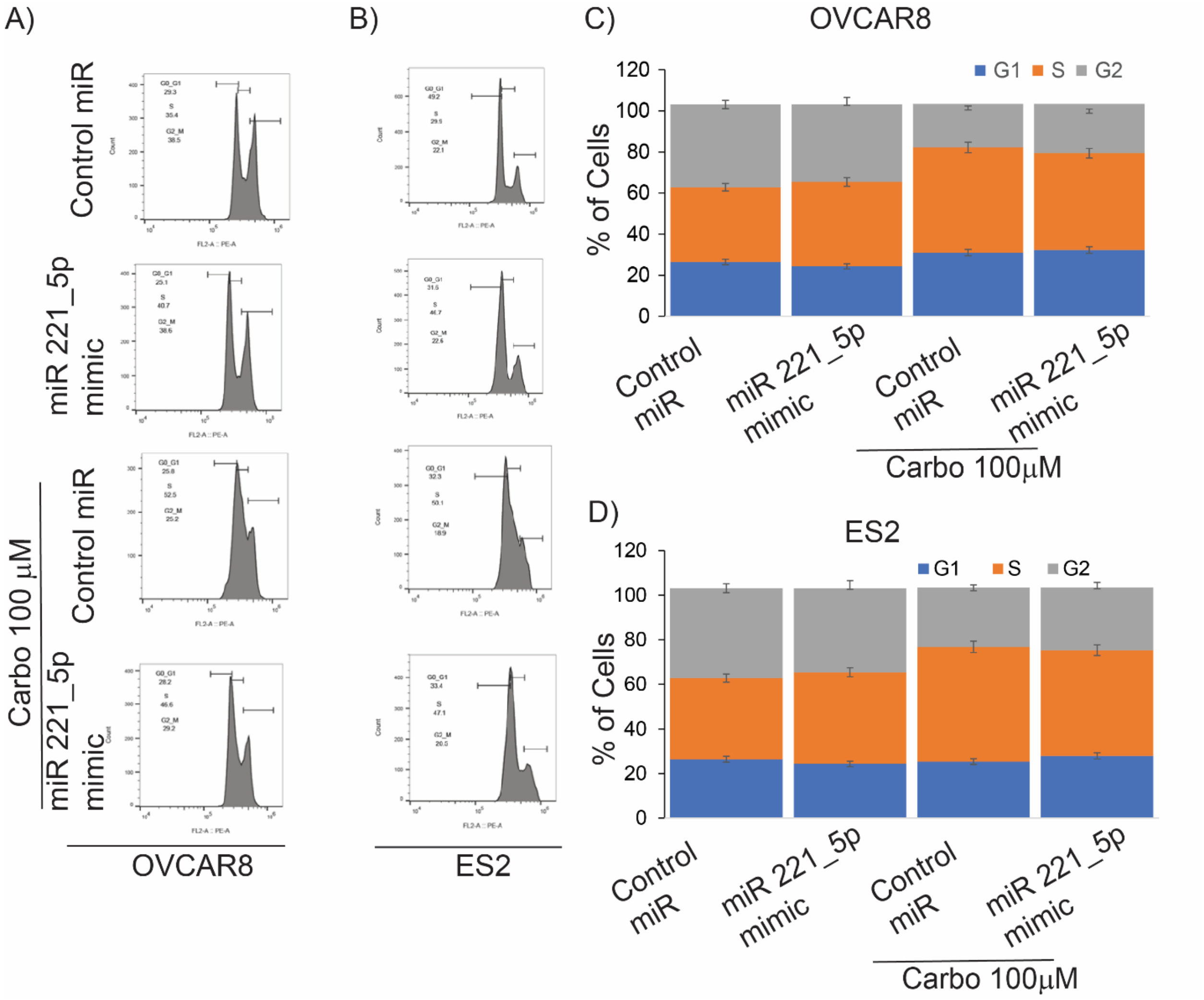


(A,B) A representative flow cytometric profile of OVCAR8 (A) and ES2 (B) cells following propidium iodide (PI) staining. Cells were transfected with Control miR or miR-221-5P mimics and, where indicated, treated with carboplatin (100 μM) for 24 h. Histograms display the DNA content and the resulting distribution of cells across the $G_0/G_1$, S and $G_2/M$ phases of the cell cycle.

(C,D) Quantitative analysis of the cell cycle distributions in OVCAR8 (C) and ES2 (D) cells 48 h post-transfection. Data represents the percentage of the total population of each phase. Data are presented as the mean ± SD of three independent biological replicates. Statistical analysis was performed using a two-tailed Student's -test (ns: non-significant).

**Supplementary Figure 5. miR-221-5p restoration depletes RAD18 and RAD51 in primary patient-derived OC cells.**

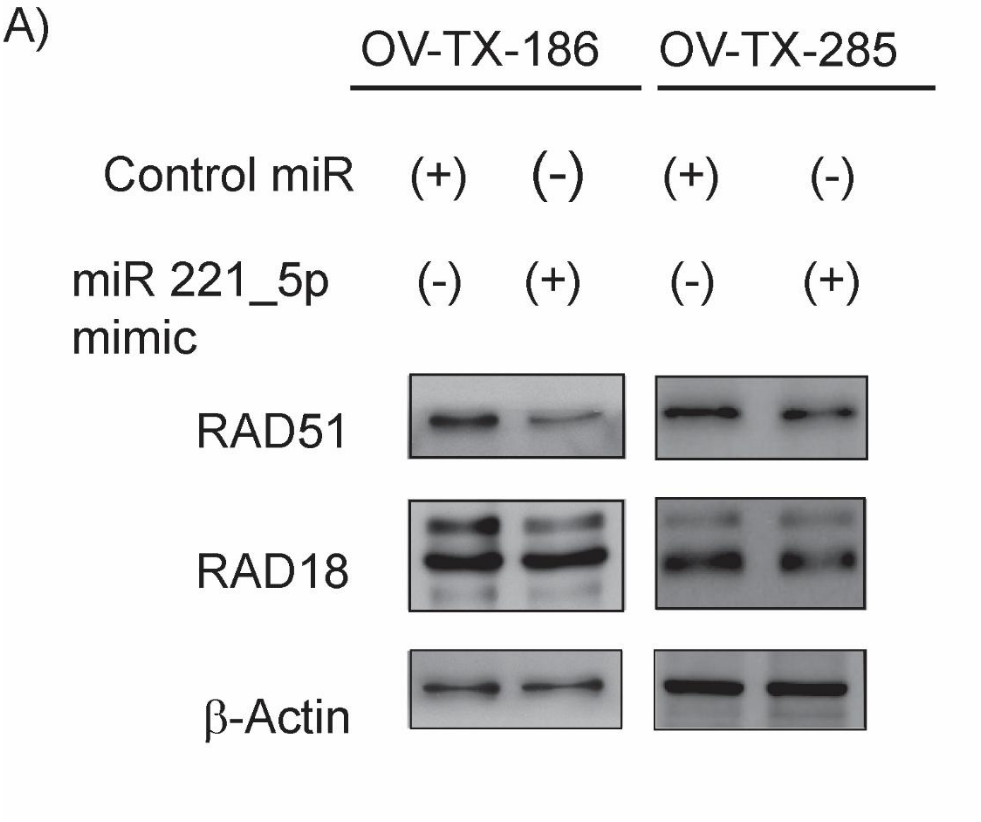


(A) Immuno blot analysis of RAD18 and RAD51 expression in two primary patient-derived ovarian cancer cell lines, OV-TX-186 and OV-TX-285, 48 h post-transfection with Control miR or miR-221-5P mimics. β-Actin and vinculin were used as loading controls to ensure equal protein distribution across lanes.

**Supplementary Figure 6. Stable miR-221-5p expression suppresses RAD18/RAD51 and attenuates 3D spheroid growth in SKOV3-Luc cells.**

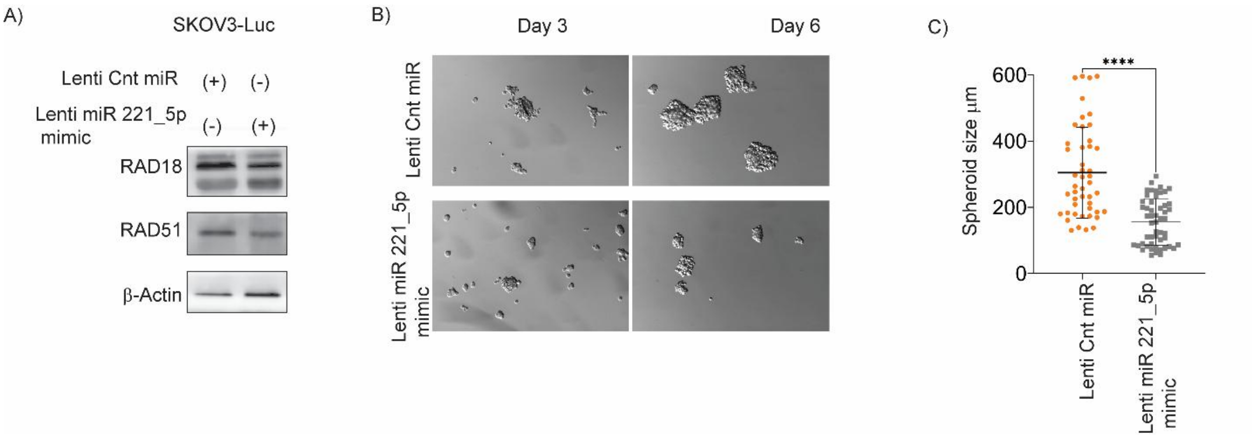


(A) Immunoblot analysis of RAD18 and RAD51 in SKOV3-Luc cells stably transduced with a control lentiviral miRNA (Lenti-Ctrl-miR) or a miR-221-5p mimic. β-Actin serves as a loading control.

(B) Representative brightfield images of 3D spheroids generated from SKOV3-Luc cells expressing control or miR-221-5p lentiviral constructs. Images illustrate the reduction in spheroid assembly and compact growth upon stable miR-221-5p expression. (C) Quantitative analysis of spheroid size (µm) in SKOV3-luc cells stably transduced with a control lentiviral miRNA (Lenti Cnt miR) or a miR-221-5p mimic. Each data point represents an individual spheroid; horizontal lines denote mean ± SD, all the spheroids bigger than 50 µm were calculated for plotting. (****$P < 0.0001$, two-tailed *t*-test).

**Supplementary Figure 8. miR-221-5p restoration sensitizes SKOV3 cells to the PARP inhibitor olaparib.**

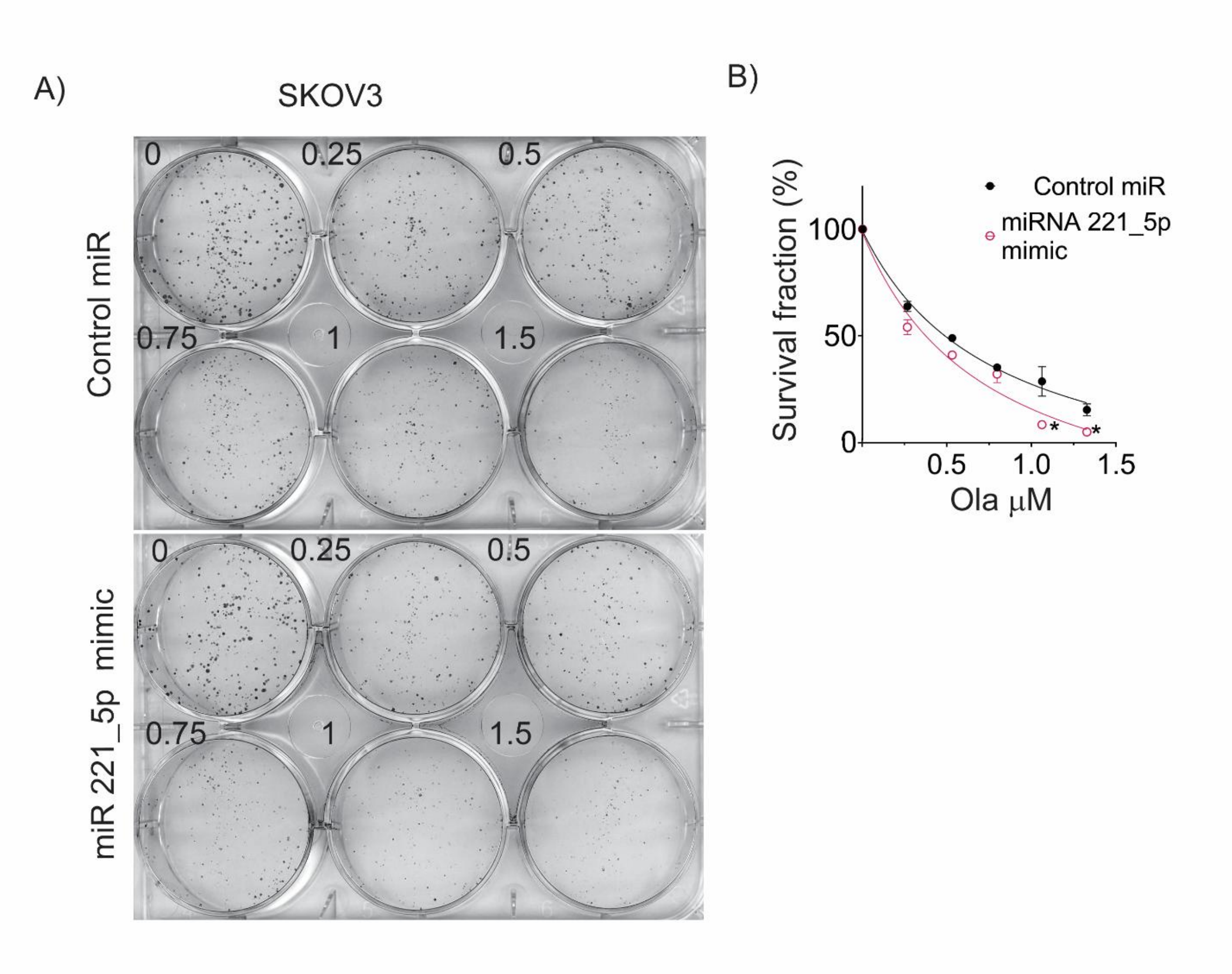


(A) Representative low-density colony formation assay using SKOV3 cells transfected with Control miR or miR-221-5P mimic and treated with increasing doses of olaparib (Ola, 0–1.5 μM). The experiment was performed in triplicates.

(B) Quantitative survival fraction analysis of SKOV3 cells transfected with Control miR or miR-221-5P mimic and treated with varying concentrations of olaparib. (*$P < 0.05$, two-tailed *t*-test).

**Table 1 : Top Significantly Differentially Expressed miRNAs Identified from GSE235525 Based on Adjusted P-Value Ranking.**

Re-analysis of the GEO dataset GSE235525 comparing serum miRNA expression profiles from HGSOC patients (CANCER) versus healthy controls (HEALTHY). Log rank values based on adjusted to P-value ranking is represented here.

Total entries: 576

| ID | log2(fold change) | -log10(Pvalue) |
|---|---|---|
| hsa-let-7b-5p | -0.761721055 | 5.593294758 |
| hsa-let-7c-5p | -0.298466886 | 2.120039421 |
| hsa-let-7d-5p | -0.459809209 | 2.072763577 |
| hsa-let-7e-5p | -0.540889674 | 4.412845267 |
| hsa-let-7g-5p | -0.795100542 | 5.928749954 |
| hsa-let-7i-5p | -0.494338187 | 2.583394954 |
| hsa-miR-1-5p | -0.554903339 | 3.454950949 |
| hsa-miR-100-5p | -0.458996509 | 2.38100408 |
| hsa-miR-101-3p | -0.548751739 | 3.781680542 |
| hsa-miR-103a-3p | -0.350425427 | 1.637249526 |
| hsa-miR-105-5p | -0.526197852 | 3.782971604 |
| hsa-miR-106b-5p | -1.053391626 | 5.168153751 |
| hsa-miR-107 | -0.844095781 | 6.846748379 |
| hsa-miR-10a-5p | -0.270305548 | 1.768111615 |
| hsa-miR-1178-3p | -0.303336182 | 2.052313933 |
| hsa-miR-1180-3p | -0.436709551 | 2.067856879 |
| hsa-miR-1185-1-3p | -0.400571747 | 1.462685329 |
| hsa-miR-1185-2-3p | -0.657495703 | 3.308508452 |
| hsa-miR-1185-5p | -0.461871813 | 3.076127602 |
| hsa-miR-1193 | -0.502102247 | 2.430886359 |
| hsa-miR-1197 | -0.33796643 | 1.890799737 |
| hsa-miR-1200 | -0.434404853 | 2.967444138 |
| hsa-miR-1202 | -0.407189729 | 2.791837397 |
| hsa-miR-1203 | -0.535338824 | 1.781732302 |
| hsa-miR-1204 | -0.338327862 | 1.95804411 |
| hsa-miR-1205 | -0.416276207 | 1.758512844 |
| hsa-miR-1206 | -0.398508852 | 2.105317179 |
| hsa-miR-122-5p | -0.559863192 | 3.286861199 |
| hsa-miR-1224-5p | -0.49059251 | 2.12034053 |
| hsa-miR-1226-3p | -0.654141109 | 3.03494588 |
| hsa-miR-1228-3p | -0.51045263 | 2.994577585 |
| hsa-miR-1233-3p | -0.305897766 | 2.758335724 |
| hsa-miR-1234-3p | -0.750483322 | 4.015567074 |
| hsa-miR-1236-3p | -0.600505547 | 3.77163568 |
| hsa-miR-124-3p | -0.426280182 | 2.423511019 |
| hsa-miR-1244 | -0.56596154 | 2.551401552 |
| hsa-miR-1245a | -0.508712333 | 2.919284007 |

| | | |
|---|---|---|
| hsa-miR-1245b-3p | -0.52072268 | 4.0996344 |
| hsa-miR-1245b-5p | -0.55371184 | 2.370413654 |
| hsa-miR-1246 | -0.505483784 | 2.859763253 |
| hsa-miR-1247-5p | 1.07487136 | 5.283803532 |
| hsa-miR-1248 | -0.767798727 | 3.719127985 |
| hsa-miR-1249-3p | -0.798594753 | 3.105581863 |
| hsa-miR-1249-5p | -0.312576938 | 1.737147942 |
| hsa-miR-1250-5p | -0.737007713 | 3.45865068 |
| hsa-miR-1252-5p | -0.623467853 | 2.735433615 |
| hsa-miR-1253 | -0.464450986 | 2.362360388 |
| hsa-miR-1255a | -0.363061204 | 1.558475555 |
| hsa-miR-1255b-5p | -0.436168626 | 2.447019213 |
| hsa-miR-125a-3p | -0.375936827 | 2.129088851 |
| hsa-miR-125a-5p | -0.363981492 | 2.018806116 |
| hsa-miR-125b-5p | -0.612700765 | 3.605018055 |
| hsa-miR-126-3p | -0.289123258 | 1.496402586 |
| hsa-miR-1260a | -1.051391387 | 6.455270905 |
| hsa-miR-1260b | -0.428022898 | 1.968696968 |
| hsa-miR-1261 | -0.589862242 | 3.053482539 |
| hsa-miR-1262 | -0.479594721 | 2.843686003 |
| hsa-miR-1266-5p | -0.484349131 | 1.913398059 |
| hsa-miR-1268b | -0.381219695 | 1.639209325 |
| hsa-miR-1269a | -0.267399585 | 1.50994415 |
| hsa-miR-1269b | -0.508803475 | 2.533962103 |
| hsa-miR-127-3p | -0.548762469 | 2.955395084 |
| hsa-miR-127-5p | -0.367095824 | 1.883680728 |
| hsa-miR-1270 | -0.744134593 | 3.857427502 |
| hsa-miR-1271-3p | -0.509534311 | 3.496372811 |
| hsa-miR-1271-5p | -1.031191711 | 3.893041403 |
| hsa-miR-1275 | -0.400962211 | 1.496708889 |
| hsa-miR-1278 | -0.455118317 | 2.33494301 |
| hsa-miR-1279 | -0.428680355 | 3.18274702 |
| hsa-miR-128-1-5p | -0.407815149 | 1.848662144 |
| hsa-miR-128-3p | -0.618225901 | 2.613590047 |
| hsa-miR-1281 | -0.460760714 | 3.325019808 |
| hsa-miR-1283 | -0.478814825 | 3.127305228 |
| hsa-miR-1285-5p | -0.384461888 | 2.075778315 |
| hsa-miR-1287-3p | -0.603925423 | 4.487269631 |
| hsa-miR-1287-5p | -0.505308714 | 2.27747446 |
| hsa-miR-1288-3p | -0.341148709 | 1.545512088 |
| hsa-miR-1289 | -0.38664985 | 1.606773697 |
| hsa-miR-129-2-3p | -0.385118243 | 1.798448127 |
| hsa-miR-129-5p | -0.492931383 | 2.207386951 |
| hsa-miR-1290 | -0.374696251 | 1.620924059 |
| hsa-miR-1296-5p | -0.313259031 | 1.777924545 |

| | | |
|---|---|---|
| hsa-miR-1297 | -0.565185188 | 2.159971767 |
| hsa-miR-1298-5p | -0.536186533 | 1.822578366 |
| hsa-miR-1301-3p | -0.492928573 | 2.856307641 |
| hsa-miR-1302 | -0.626931672 | 3.557562828 |
| hsa-miR-1303 | -0.328567683 | 1.937716798 |
| hsa-miR-1306-5p | -0.518159988 | 2.893309735 |
| hsa-miR-1307-3p | -0.612267733 | 2.409087163 |
| hsa-miR-130a-3p | -0.387437589 | 2.217110092 |
| hsa-miR-130b-3p | -0.718637785 | 3.11483014 |
| hsa-miR-1322 | -0.314957382 | 1.753598061 |
| hsa-miR-133a-3p | -0.452706595 | 3.098210118 |
| hsa-miR-133a-5p | -0.493438725 | 3.390298362 |
| hsa-miR-133b | -0.58866441 | 3.539114796 |
| hsa-miR-134-3p | -0.669905223 | 2.934897641 |
| hsa-miR-135b-5p | -0.483353098 | 1.84041785 |
| hsa-miR-136-5p | -0.416464766 | 2.576633819 |
| hsa-miR-137 | -0.790483756 | 5.009622026 |
| hsa-miR-139-5p | -0.456517846 | 2.005188723 |
| hsa-miR-140-5p | -0.416951862 | 1.897216449 |
| hsa-miR-142-5p | -1.251468588 | 7.475796176 |
| hsa-miR-145-5p | -1.673404538 | 7.487428375 |
| hsa-miR-1469 | -0.586053775 | 3.635636866 |
| hsa-miR-146b-3p | -0.638357014 | 3.706644723 |
| hsa-miR-146b-5p | -0.40262681 | 1.868312068 |
| hsa-miR-147a | -0.548353109 | 3.935993044 |
| hsa-miR-148a-3p | -0.341463358 | 1.629523897 |
| hsa-miR-149-5p | -0.4876277 | 3.138818954 |
| hsa-miR-151a-3p | -1.204389535 | 9.655057547 |
| hsa-miR-151a-5p | -0.870503347 | 2.776246228 |
| hsa-miR-151b | -0.81859878 | 3.937295076 |
| hsa-miR-152-3p | -0.439921975 | 2.329209846 |
| hsa-miR-152-5p | -0.583605841 | 3.060842517 |
| hsa-miR-153-3p | -0.534371586 | 2.209712869 |
| hsa-miR-1537-3p | -0.519369244 | 2.900198113 |
| hsa-miR-154-5p | -0.378350112 | 1.750893922 |
| hsa-miR-155-5p | -0.445244147 | 1.77657908 |
| hsa-miR-15b-5p | -1.149923819 | 7.919275058 |
| hsa-miR-16-5p | -1.077516767 | 8.220750236 |
| hsa-miR-181a-2-3p | -1.134818552 | 3.493963224 |
| hsa-miR-181a-3p | -0.437135628 | 3.004602974 |
| hsa-miR-181b-2-3p | -0.704889337 | 4.295694396 |
| hsa-miR-181c-5p | -0.391971092 | 1.739559055 |
| hsa-miR-181d-3p | -0.421451522 | 2.178854244 |
| hsa-miR-182-3p | -0.302048506 | 1.673031133 |
| hsa-miR-182-5p | -0.380177328 | 2.125712016 |

| | | |
|---|---|---|
| hsa-miR-1827 | -0.540457311 | 1.733617993 |
| hsa-miR-184 | -0.384005221 | 2.805774108 |
| hsa-miR-185-5p | -0.61885273 | 3.705031227 |
| hsa-miR-186-5p | -0.715353086 | 4.980099289 |
| hsa-miR-187-3p | -0.261602942 | 1.627039088 |
| hsa-miR-188-3p | -0.398354136 | 2.588941741 |
| hsa-miR-188-5p | -0.481855163 | 2.479580119 |
| hsa-miR-18a-5p | -0.379585177 | 2.383434387 |
| hsa-miR-18b-5p | -0.439851674 | 3.30181487 |
| hsa-miR-1908-5p | -0.516967848 | 1.97782087 |
| hsa-miR-190a-3p | -0.367522034 | 1.717949585 |
| hsa-miR-190a-5p | -0.308807053 | 1.778572891 |
| hsa-miR-190b | -0.567801909 | 4.038786978 |
| hsa-miR-1910-3p | -0.900471332 | 6.765223786 |
| hsa-miR-1910-5p | -0.39421091 | 1.762704185 |
| hsa-miR-1915-3p | -0.569787411 | 3.607806789 |
| hsa-miR-193a-5p+hsa-miR-193b-5p | -0.639893304 | 3.687080646 |
| hsa-miR-194-5p | -0.556008336 | 2.767165401 |
| hsa-miR-195-5p | -0.452130236 | 2.163440242 |
| hsa-miR-196a-3p | -0.458650685 | 2.920343776 |
| hsa-miR-197-3p | -0.612928739 | 3.033121033 |
| hsa-miR-197-5p | -0.461854446 | 2.530642244 |
| hsa-miR-1972 | -0.257593148 | 1.531377473 |
| hsa-miR-1976 | -0.401795809 | 3.046812763 |
| hsa-miR-198 | -0.382039928 | 2.076494752 |
| hsa-miR-199a-3p+hsa-miR-199b-3p | -0.35237602 | 1.852601743 |
| hsa-miR-199a-5p | -0.706858528 | 5.606357212 |
| hsa-miR-199b-5p | -0.620573002 | 4.875113997 |
| hsa-miR-19b-3p | -0.704185774 | 3.545628331 |
| hsa-miR-200a-3p | -0.687262833 | 2.866422657 |
| hsa-miR-200b-3p | -0.390133264 | 2.686824753 |
| hsa-miR-200c-3p | -0.434488781 | 1.857628791 |
| hsa-miR-203a-3p | -0.513498411 | 2.340923441 |
| hsa-miR-204-5p | -0.360784626 | 2.787703546 |
| hsa-miR-206 | -0.372979158 | 1.723918549 |
| hsa-miR-208a-3p | -0.335279104 | 1.701219055 |
| hsa-miR-208b-3p | -0.309831937 | 1.648531837 |
| hsa-miR-20a-5p+hsa-miR-20b-5p | -0.535831574 | 2.265330111 |
| hsa-miR-21-5p | -0.902705183 | 4.147039987 |
| hsa-miR-211-5p | -0.560463964 | 2.115331227 |
| hsa-miR-2110 | -0.40394887 | 2.178777573 |
| hsa-miR-2116-5p | -0.466042547 | 3.11579981 |

| | | |
|---|---|---|
| hsa-miR-2117 | -0.508966074 | 2.445449104 |
| hsa-miR-214-3p | -0.530190705 | 5.058706381 |
| hsa-miR-216a-5p | -0.322198433 | 1.551773312 |
| hsa-miR-216b-5p | -0.477617042 | 3.279651126 |
| hsa-miR-217 | -0.475562998 | 2.327789771 |
| hsa-miR-218-5p | -0.342172892 | 2.190948959 |
| hsa-miR-219a-5p | -0.531797843 | 2.275839998 |
| hsa-miR-219b-3p | -0.748014454 | 3.671223378 |
| hsa-miR-22-3p | -0.515328761 | 2.042754686 |
| hsa-miR-221-5p | -0.915760956 | 5.696653297 |
| hsa-miR-222-3p | -0.46684721 | 1.830598638 |
| hsa-miR-224-5p | -0.513761147 | 1.564360823 |
| hsa-miR-2278 | -0.492030753 | 2.894562197 |
| hsa-miR-23a-3p | -0.579180811 | 2.120626459 |
| hsa-miR-23c | -0.313980463 | 2.002570062 |
| hsa-miR-24-3p | -0.46491602 | 3.139877441 |
| hsa-miR-25-3p | -0.686683266 | 2.912403223 |
| hsa-miR-25-5p | -0.533879078 | 2.192231517 |
| hsa-miR-2682-5p | -0.256123968 | 1.601416862 |
| hsa-miR-26b-5p | -0.35062096 | 1.998867943 |
| hsa-miR-27a-3p | -0.68791203 | 5.143492571 |
| hsa-miR-27b-3p | -0.37345265 | 2.411982136 |
| hsa-miR-28-3p | -0.532737305 | 3.438817916 |
| hsa-miR-28-5p | -0.615947942 | 2.75818421 |
| hsa-miR-296-3p | -0.599914152 | 6.044430714 |
| hsa-miR-296-5p | -0.404919593 | 2.084582748 |
| hsa-miR-297 | -0.527109466 | 3.555591208 |
| hsa-miR-298 | -0.570995417 | 1.797589837 |
| hsa-miR-299-3p | -0.378485202 | 2.356285823 |
| hsa-miR-299-5p | -0.489436817 | 3.873636948 |
| hsa-miR-29a-3p | -0.295574131 | 1.913736422 |
| hsa-miR-29c-3p | -0.675684744 | 4.110931587 |
| hsa-miR-300 | -0.547505415 | 3.851922102 |
| hsa-miR-301a-3p | -0.335523869 | 1.548718195 |
| hsa-miR-301a-5p | -0.347032094 | 2.327709259 |
| hsa-miR-301b-3p | -0.254663062 | 1.5812709 |
| hsa-miR-301b-5p | -0.30216586 | 1.940392481 |
| hsa-miR-302a-5p | -0.503975315 | 3.956530655 |
| hsa-miR-302c-3p | -0.508710917 | 2.296414354 |
| hsa-miR-3065-3p | -0.338724505 | 1.509923148 |
| hsa-miR-3065-5p | -0.307566642 | 1.66127882 |
| hsa-miR-3074-3p | -0.456655877 | 1.790265548 |
| hsa-miR-30a-3p | -0.704205411 | 3.249191867 |
| hsa-miR-30c-5p | -0.567622031 | 3.601933978 |
| hsa-miR-30d-5p | -0.533457237 | 2.368516936 |

| | | |
|---|---|---|
| hsa-miR-30e-5p | -0.287385502 | 1.599835271 |
| hsa-miR-3127-5p | -0.563546422 | 2.481234135 |
| hsa-miR-3130-3p | -0.458850991 | 2.058257695 |
| hsa-miR-3131 | -0.892830055 | 2.360915207 |
| hsa-miR-3136-5p | -0.316097884 | 1.481857508 |
| hsa-miR-3140-3p | -0.383637603 | 2.033677842 |
| hsa-miR-3144-5p | -0.29349314 | 2.518232312 |
| hsa-miR-3147 | -0.373909591 | 1.738472397 |
| hsa-miR-3150b-3p | -0.399218727 | 3.366184674 |
| hsa-miR-3161 | -0.516851311 | 2.366429725 |
| hsa-miR-3168 | -0.635805423 | 3.768031155 |
| hsa-miR-3179 | -0.260908448 | 1.487136234 |
| hsa-miR-3180-3p | -0.610913992 | 2.544798207 |
| hsa-miR-3180-5p | -0.579322197 | 3.232854894 |
| hsa-miR-3185 | -0.414342534 | 1.926928197 |
| hsa-miR-3190-3p | -0.50321052 | 2.630827674 |
| hsa-miR-3192-5p | -0.561746818 | 2.475474863 |
| hsa-miR-3195 | -0.429715638 | 1.496821272 |
| hsa-miR-32-5p | -0.494699103 | 2.517935019 |
| hsa-miR-3202 | -0.845015607 | 6.748011871 |
| hsa-miR-320a | -0.455713383 | 1.991481799 |
| hsa-miR-320b | -0.336325728 | 1.944887113 |
| hsa-miR-320c | -0.34355085 | 1.590800079 |
| hsa-miR-320d | -0.645559722 | 4.317109771 |
| hsa-miR-323b-5p | -0.438207744 | 2.661591475 |
| hsa-miR-324-5p | -0.357115465 | 2.44125155 |
| hsa-miR-325 | -0.682952272 | 2.798886333 |
| hsa-miR-326 | -0.50268513 | 2.052434875 |
| hsa-miR-328-3p | -0.520836481 | 2.306153542 |
| hsa-miR-328-5p | -0.513259035 | 1.650696201 |
| hsa-miR-329-5p | -0.472082278 | 1.619356341 |
| hsa-miR-330-3p | -0.383827803 | 1.467073409 |
| hsa-miR-330-5p | -0.497556132 | 2.385749737 |
| hsa-miR-331-3p | -0.502278555 | 2.20134136 |
| hsa-miR-331-5p | -0.51912319 | 2.448175716 |
| hsa-miR-335-5p | -0.546194911 | 2.119074259 |
| hsa-miR-337-3p | -0.826609057 | 3.297024773 |
| hsa-miR-337-5p | -0.440785642 | 3.500910805 |
| hsa-miR-339-3p | -0.307861147 | 1.515565669 |
| hsa-miR-339-5p | -0.333611824 | 1.502347549 |
| hsa-miR-33a-5p | -0.394995661 | 2.743601253 |
| hsa-miR-33b-5p | -0.953379504 | 3.653179046 |
| hsa-miR-340-5p | -0.294646929 | 1.891061985 |
| hsa-miR-342-3p | -0.725582224 | 4.849462312 |
| hsa-miR-342-5p | -0.704587477 | 4.686885631 |

| | | |
|---|---|---|
| hsa-miR-345-3p | -0.415316646 | 1.983660028 |
| hsa-miR-345-5p | -0.570736788 | 4.320207401 |
| hsa-miR-346 | -0.456938753 | 1.766150108 |
| hsa-miR-34a-5p | -0.497228886 | 3.082680464 |
| hsa-miR-34c-5p | -0.85198168 | 2.58694049 |
| hsa-miR-3605-5p | -0.450462267 | 2.629624657 |
| hsa-miR-361-5p | -0.375434236 | 2.075358646 |
| hsa-miR-3613-5p | -0.347171274 | 2.008304995 |
| hsa-miR-3614-3p | -0.370810092 | 1.639094677 |
| hsa-miR-3614-5p | -0.486061311 | 1.710255547 |
| hsa-miR-3615 | -0.461254297 | 3.31002617 |
| hsa-miR-362-5p | -0.196336411 | 1.607604633 |
| hsa-miR-363-5p | -0.624459785 | 4.040378733 |
| hsa-miR-365a-3p+hsa-miR-365b-3p | -0.351863335 | 1.639331595 |
| hsa-miR-367-3p | -0.571474964 | 3.122811911 |
| hsa-miR-369-3p | -0.489732184 | 2.036300642 |
| hsa-miR-369-5p | -0.401570473 | 1.905085017 |
| hsa-miR-3690 | -0.414442969 | 1.77326278 |
| hsa-miR-370-3p | -0.566340482 | 2.107233637 |
| hsa-miR-371a-5p | -0.472329072 | 2.55912679 |
| hsa-miR-373-3p | -0.507049099 | 2.796578395 |
| hsa-miR-374b-5p | -0.769517823 | 4.973956088 |
| hsa-miR-374c-5p | -0.679829292 | 4.140469348 |
| hsa-miR-375 | -0.428869136 | 1.740356951 |
| hsa-miR-376a-2-5p | -0.46707961 | 2.083912219 |
| hsa-miR-376a-3p | -0.48426439 | 2.033977135 |
| hsa-miR-376b-3p | -0.500643674 | 2.150730387 |
| hsa-miR-376c-3p | -0.619393751 | 3.261434687 |
| hsa-miR-376c-5p | -0.901843017 | 3.510258401 |
| hsa-miR-377-3p | -0.403660717 | 1.647215532 |
| hsa-miR-378c | -0.345772208 | 1.590844099 |
| hsa-miR-378e | -0.478427328 | 4.08621545 |
| hsa-miR-378g | -0.352817329 | 2.93466307 |
| hsa-miR-378i | -0.383504296 | 2.823380331 |
| hsa-miR-379-5p | -0.442340235 | 2.977863282 |
| hsa-miR-381-5p | -0.401950962 | 2.087809824 |
| hsa-miR-382-3p | -0.569159591 | 3.886506096 |
| hsa-miR-383-5p | -0.453722007 | 2.380566218 |
| hsa-miR-384 | -0.39695019 | 1.557850508 |
| hsa-miR-3916 | -0.422839999 | 1.869765051 |
| hsa-miR-3918 | -0.469417528 | 3.364409108 |
| hsa-miR-3928-3p | -0.620314126 | 2.933758501 |
| hsa-miR-3934-5p | -0.728488292 | 3.098839616 |
| hsa-miR-409-5p | -0.615969308 | 3.227355055 |

| | | |
|---|---|---|
| hsa-miR-410-3p | -0.560156671 | 1.973955463 |
| hsa-miR-411-5p | -0.408676846 | 1.896188595 |
| hsa-miR-421 | -0.419625293 | 3.792265786 |
| hsa-miR-422a | -0.368215164 | 2.282195559 |
| hsa-miR-423-3p | -0.563121442 | 2.852583151 |
| hsa-miR-424-5p | -0.519671167 | 3.746445659 |
| hsa-miR-425-5p | -0.37302547 | 2.531945226 |
| hsa-miR-4284 | -0.986084789 | 7.30341204 |
| hsa-miR-4286 | -0.467056047 | 4.472113106 |
| hsa-miR-429 | -0.315605917 | 2.210845011 |
| hsa-miR-431-5p | -0.74916779 | 3.360201754 |
| hsa-miR-432-5p | -0.425357987 | 1.73243104 |
| hsa-miR-433-3p | -0.493114016 | 2.46293953 |
| hsa-miR-433-5p | -0.346413909 | 1.688387268 |
| hsa-miR-4425 | -0.554294004 | 3.369408976 |
| hsa-miR-4431 | -0.565583401 | 2.340324809 |
| hsa-miR-4443 | -0.58453765 | 3.705332758 |
| hsa-miR-4448 | -0.946297954 | 3.256333059 |
| hsa-miR-4451 | -0.410071945 | 1.644595328 |
| hsa-miR-4454+hsa-miR-7975 | -0.498988651 | 3.035169002 |
| hsa-miR-4455 | 0.96820289 | 3.336343985 |
| hsa-miR-4458 | -0.611519545 | 2.253871962 |
| hsa-miR-4461 | -0.461890946 | 1.570132599 |
| hsa-miR-448 | -0.309323984 | 1.719929898 |
| hsa-miR-4485-3p | -0.42465525 | 2.440311233 |
| hsa-miR-4488 | -0.460572468 | 3.289868668 |
| hsa-miR-449a | -0.469128642 | 2.76828538 |
| hsa-miR-449c-5p | -0.267727195 | 1.520891761 |
| hsa-miR-450a-1-3p | -0.501686584 | 2.713794502 |
| hsa-miR-450a-2-3p | -0.497003923 | 1.928877913 |
| hsa-miR-450b-3p | -0.289487973 | 1.472311517 |
| hsa-miR-450b-5p | -0.404685084 | 2.219659229 |
| hsa-miR-4516 | -0.41475129 | 1.749339802 |
| hsa-miR-452-5p | -1.432531152 | 4.357444818 |
| hsa-miR-4524a-5p | -0.484056429 | 2.579914617 |
| hsa-miR-4532 | -0.521627125 | 2.073969244 |
| hsa-miR-4536-5p | -0.451702705 | 3.404397052 |
| hsa-miR-455-3p | -0.540698579 | 4.280541508 |
| hsa-miR-4707-5p | -0.385070141 | 1.501650264 |
| hsa-miR-4741 | -0.272518096 | 1.718153198 |
| hsa-miR-4755-5p | -0.395313332 | 2.003279905 |
| hsa-miR-4787-3p | -0.474277672 | 2.44562072 |
| hsa-miR-4787-5p | -0.488853723 | 2.067491956 |
| hsa-miR-483-3p | -0.399736654 | 2.252973353 |

| | | |
|---|---|---|
| hsa-miR-483-5p | -0.425764073 | 1.582203913 |
| hsa-miR-485-3p | -0.459126956 | 3.016877564 |
| hsa-miR-486-3p | -0.645092352 | 3.729706088 |
| hsa-miR-487a-3p | -0.616017687 | 5.467037458 |
| hsa-miR-487b-3p | -0.43447419 | 3.144660191 |
| hsa-miR-487b-5p | -0.35433106 | 1.93376392 |
| hsa-miR-488-3p | -0.371615263 | 2.278783524 |
| hsa-miR-490-5p | -0.593137597 | 4.185851429 |
| hsa-miR-491-5p | -0.568630873 | 2.849146807 |
| hsa-miR-492 | -0.556625827 | 2.881915758 |
| hsa-miR-493-3p | -0.479202916 | 2.220728787 |
| hsa-miR-494-3p | -0.426407974 | 2.417432753 |
| hsa-miR-495-5p | -0.756956418 | 5.996531432 |
| hsa-miR-496 | -0.375943491 | 2.878836299 |
| hsa-miR-498 | -1.253904904 | 3.365790493 |
| hsa-miR-499b-3p | -0.527571743 | 3.068080045 |
| hsa-miR-499b-5p | -0.521328849 | 3.715888398 |
| hsa-miR-500a-5p+hsa-miR-501-5p | -0.499071973 | 3.984248186 |
| hsa-miR-501-3p | -0.585568222 | 2.943475355 |
| hsa-miR-5010-3p | -0.733874746 | 3.091119386 |
| hsa-miR-5010-5p | -0.538870474 | 3.812399449 |
| hsa-miR-502-3p | -0.545947209 | 1.808884206 |
| hsa-miR-502-5p | -0.344631577 | 1.894942368 |
| hsa-miR-503-5p | -0.41373189 | 2.212916253 |
| hsa-miR-504-5p | -0.517535661 | 1.670263411 |
| hsa-miR-505-3p | -1.194743075 | 3.687847909 |
| hsa-miR-506-5p | -0.345961363 | 2.261004821 |
| hsa-miR-508-5p | -0.333664112 | 1.463054878 |
| hsa-miR-509-3-5p | -0.32574091 | 2.100847372 |
| hsa-miR-509-3p | -0.342149141 | 1.569905486 |
| hsa-miR-509-5p | -0.537107033 | 3.013722219 |
| hsa-miR-510-5p | -0.401427454 | 3.001285441 |
| hsa-miR-511-5p | -0.754505507 | 4.865659481 |
| hsa-miR-512-3p | -0.316395276 | 1.525007696 |
| hsa-miR-512-5p | -0.511345826 | 1.776457587 |
| hsa-miR-513a-3p | -0.495349723 | 4.162944946 |
| hsa-miR-513a-5p | -0.4649856 | 3.628407019 |
| hsa-miR-513c-3p | -0.560941186 | 3.003628615 |
| hsa-miR-514a-3p | -0.560280445 | 3.312146721 |
| hsa-miR-514a-5p | -0.268841868 | 1.651297112 |
| hsa-miR-514b-3p | -0.662945165 | 3.916459431 |
| hsa-miR-515-3p | -0.324382702 | 2.188526238 |
| hsa-miR-515-5p | -0.390705065 | 2.187230557 |

| | | |
|---|---|---|
| hsa-miR-516a-3p+hsa-miR-516b-3p | -0.423253446 | 1.629972688 |
| hsa-miR-517a-3p | -0.673329934 | 3.800051961 |
| hsa-miR-517b-3p | -0.563782653 | 4.272048951 |
| hsa-miR-518c-3p | -0.368405962 | 1.651094157 |
| hsa-miR-5196-3p+hsa-miR-6732-3p | -0.439232801 | 2.748968783 |
| hsa-miR-5196-5p | -0.442096843 | 1.546256346 |
| hsa-miR-519b-5p+hsa-miR-519c-5p+hsa-miR-523-5p+hsa-miR-518e-5p+hsa-miR-522-5p+hsa-miR-519a-5p | -0.516652557 | 2.449205021 |
| hsa-miR-519c-3p | -0.768750593 | 2.71386341 |
| hsa-miR-519d-3p | -0.292545083 | 1.788375552 |
| hsa-miR-519e-3p | -0.370609474 | 1.998668815 |
| hsa-miR-520a-3p | -0.772515813 | 4.031607719 |
| hsa-miR-520b | -0.391717084 | 1.999699584 |
| hsa-miR-520c-3p | -0.543429995 | 2.756104283 |
| hsa-miR-520d-3p | -0.41654738 | 2.722377408 |
| hsa-miR-520d-5p+hsa-miR-527+hsa-miR-518a-5p | -0.515417064 | 2.368154183 |
| hsa-miR-520e | -0.489903949 | 1.881090988 |
| hsa-miR-520g-3p | -0.561454127 | 2.009803652 |
| hsa-miR-520h | -0.407360393 | 1.77249714 |
| hsa-miR-521 | -0.430217495 | 1.789126696 |
| hsa-miR-523-3p | -0.510769173 | 2.975099354 |
| hsa-miR-524-3p | -0.575503014 | 1.82230927 |
| hsa-miR-525-3p | -0.697183555 | 3.128168466 |
| hsa-miR-525-5p | -0.871635533 | 2.325935856 |
| hsa-miR-526a+hsa-miR-518c-5p+hsa-miR-518d-5p | -0.366773667 | 2.762559582 |
| hsa-miR-532-3p | -0.435793061 | 2.910187535 |
| hsa-miR-539-3p | -0.522298594 | 1.905271177 |
| hsa-miR-541-3p | -0.415319398 | 1.735749994 |
| hsa-miR-542-5p | -0.318786115 | 1.813215049 |
| hsa-miR-544a | -0.290195576 | 2.376894171 |
| hsa-miR-548a-3p | -0.451208142 | 2.263479733 |
| hsa-miR-548a-5p | -0.551292805 | 2.875508962 |
| hsa-miR-548aa+hsa-miR-548t-3p | -0.293325431 | 1.925525745 |
| hsa-miR-548ad-3p | -0.590718134 | 1.577689668 |
| hsa-miR-548ah-5p | -0.33527028 | 1.950796568 |

| | | |
|---|---|---|
| hsa-miR-548ai+hsa-miR-570-5p | -0.381424267 | 2.356768431 |
| hsa-miR-548ak | -0.589502302 | 2.188110559 |
| hsa-miR-548ar-3p | -0.561939302 | 3.035513419 |
| hsa-miR-548ar-5p | -0.369668281 | 2.841418484 |
| hsa-miR-548b-3p | -0.277819064 | 1.610278752 |
| hsa-miR-548c-5p+hsa-miR-548o-5p+hsa-miR-548am-5p | -0.648399349 | 3.637711248 |
| hsa-miR-548e-3p | -0.610316185 | 2.365044711 |
| hsa-miR-548e-5p | -0.502580042 | 2.193973094 |
| hsa-miR-548h-5p | -0.436194815 | 2.557006747 |
| hsa-miR-548i | -0.539771834 | 2.373829508 |
| hsa-miR-548j-3p | -0.517746997 | 2.960389788 |
| hsa-miR-548j-5p | -0.817811335 | 2.80389766 |
| hsa-miR-548l | -0.402806774 | 2.659796628 |
| hsa-miR-548n | -0.284825024 | 1.556152058 |
| hsa-miR-548o-3p+hsa-miR-548ah-3p+hsa-miR-548av-3p | -0.874880191 | 2.489567196 |
| hsa-miR-548q | -0.719004895 | 3.879487185 |
| hsa-miR-548y | -0.580346032 | 4.489715396 |
| hsa-miR-548z+hsa-miR-548h-3p | -0.343470557 | 2.188527541 |
| hsa-miR-549a | -0.6297554 | 3.116167619 |
| hsa-miR-550a-5p | -0.51086138 | 1.54083514 |
| hsa-miR-551a | -0.47546162 | 1.787175326 |
| hsa-miR-551b-3p | -0.489726453 | 2.030028497 |
| hsa-miR-552-3p | -0.310877917 | 2.211393586 |
| hsa-miR-553 | -0.364719953 | 1.76890618 |
| hsa-miR-554 | -0.351557819 | 1.600573182 |
| hsa-miR-561-3p | -0.345563485 | 2.0939931 |
| hsa-miR-563 | -0.575859969 | 2.332880928 |
| hsa-miR-564 | -0.347959796 | 2.17565412 |
| hsa-miR-566 | -0.393060019 | 2.384255992 |
| hsa-miR-567 | -0.393489509 | 1.902382614 |
| hsa-miR-568 | -0.481972964 | 2.410499633 |
| hsa-miR-571 | -0.389579223 | 1.809872426 |
| hsa-miR-572 | -0.440648001 | 1.853893422 |
| hsa-miR-573 | -0.355441585 | 2.123803006 |
| hsa-miR-574-3p | -0.49130477 | 2.488932867 |
| hsa-miR-574-5p | -0.465139339 | 3.520929319 |
| hsa-miR-576-5p | -0.283890135 | 1.744920413 |
| hsa-miR-577 | -0.888329384 | 3.84410742 |
| hsa-miR-579-3p | -0.41834622 | 2.390629357 |

| | | |
|---|---|---|
| hsa-miR-580-3p | -0.576139799 | 2.058265726 |
| hsa-miR-582-5p | -0.352772063 | 1.967038965 |
| hsa-miR-584-3p | -0.321386018 | 2.421081376 |
| hsa-miR-584-5p | -0.26707918 | 1.739596695 |
| hsa-miR-585-3p | -0.487659383 | 2.757142003 |
| hsa-miR-587 | -0.609372478 | 2.905395867 |
| hsa-miR-589-5p | -0.294084082 | 1.487689642 |
| hsa-miR-590-5p | -0.514971091 | 2.718195216 |
| hsa-miR-591 | -0.462827878 | 1.467669983 |
| hsa-miR-593-3p | -0.452811618 | 1.949509718 |
| hsa-miR-595 | -0.427860717 | 2.094914222 |
| hsa-miR-598-3p | -0.344739084 | 2.416406606 |
| hsa-miR-599 | -0.738333627 | 4.757196707 |
| hsa-miR-600 | -0.352223259 | 1.796578841 |
| hsa-miR-601 | -0.475396255 | 2.02281931 |
| hsa-miR-604 | -0.665437103 | 2.513714255 |
| hsa-miR-607 | -0.835841431 | 4.13204709 |
| hsa-miR-610 | -0.434615977 | 2.499499812 |
| hsa-miR-615-3p | -0.40510915 | 2.927818709 |
| hsa-miR-617 | -0.42558485 | 1.999632431 |
| hsa-miR-619-3p | -0.548881393 | 2.251877241 |
| hsa-miR-620 | -0.665843032 | 3.801527006 |
| hsa-miR-624-3p | -0.317883484 | 1.483454509 |
| hsa-miR-625-5p | -0.45741156 | 2.116097788 |
| hsa-miR-627-3p | -0.458328431 | 3.136681005 |
| hsa-miR-627-5p | -0.35933425 | 2.107269897 |
| hsa-miR-628-3p | -0.693936509 | 3.252524086 |
| hsa-miR-629-5p | -0.483227329 | 2.881043445 |
| hsa-miR-630 | -0.357539738 | 1.817123153 |
| hsa-miR-637 | -0.434555513 | 2.041429181 |
| hsa-miR-639 | -0.459439932 | 1.649444259 |
| hsa-miR-640 | -0.322823484 | 2.129432216 |
| hsa-miR-642a-5p | -0.695640048 | 3.393242185 |
| hsa-miR-643 | -0.437789823 | 2.875770019 |
| hsa-miR-648 | -0.485013142 | 1.823955266 |
| hsa-miR-649 | -0.423407888 | 2.854993554 |
| hsa-miR-6503-3p | -0.428473991 | 3.397889463 |
| hsa-miR-6503-5p | -0.743765552 | 2.318848267 |
| hsa-miR-6511a-3p | -0.480812761 | 2.571492487 |
| hsa-miR-654-3p | -0.368703632 | 1.618609518 |
| hsa-miR-654-5p | -0.739430631 | 3.012285518 |
| hsa-miR-655-3p | -0.276696462 | 1.553071217 |
| hsa-miR-660-5p | -0.532282581 | 4.162206609 |
| hsa-miR-661 | -0.309766912 | 2.166913764 |
| hsa-miR-663a | -0.475005762 | 3.475127649 |

| | | |
|---|---|---|
| hsa-miR-664a-3p | -0.505678686 | 3.76420382 |
| hsa-miR-664b-3p | -0.365383658 | 2.442869031 |
| hsa-miR-664b-5p | -0.657666386 | 3.199088572 |
| hsa-miR-665 | -0.451691678 | 2.157425776 |
| hsa-miR-671-5p | -0.834464868 | 2.917772066 |
| hsa-miR-6720-3p | -0.685807289 | 2.365799165 |
| hsa-miR-6721-5p | -0.517256915 | 3.549566877 |
| hsa-miR-6724-5p | -0.721185364 | 3.196214882 |
| hsa-miR-675-5p | -0.451576244 | 2.060016352 |
| hsa-miR-7-5p | -0.357712342 | 2.035852154 |
| hsa-miR-708-5p | -0.345725852 | 2.066993988 |
| hsa-miR-758-3p+hsa-miR-411-3p | -0.69291845 | 3.344538855 |
| hsa-miR-758-5p | -0.306217277 | 1.583734943 |
| hsa-miR-760 | -1.028583968 | 4.036414478 |
| hsa-miR-764 | -0.683143527 | 3.1635574 |
| hsa-miR-766-3p | -0.473999316 | 2.982440911 |
| hsa-miR-766-5p | -0.538590416 | 2.159301483 |
| hsa-miR-767-5p | -0.386362706 | 1.644253939 |
| hsa-miR-769-3p | -0.303083083 | 1.688108065 |
| hsa-miR-769-5p | -0.388087053 | 2.457741193 |
| hsa-miR-770-5p | -0.46584854 | 2.817619877 |
| hsa-miR-802 | -0.53691187 | 3.148693183 |
| hsa-miR-873-3p | -0.356845468 | 2.354755456 |
| hsa-miR-873-5p | -0.556659099 | 1.997540799 |
| hsa-miR-874-3p | -0.61793203 | 3.281744642 |
| hsa-miR-875-3p | -0.411352232 | 2.371200587 |
| hsa-miR-876-5p | -0.370458634 | 1.921810703 |
| hsa-miR-877-5p | -0.407117855 | 2.044210372 |
| hsa-miR-885-5p | -0.357063133 | 1.584739554 |
| hsa-miR-887-3p | -0.282025881 | 1.569354594 |
| hsa-miR-887-5p | -0.392796844 | 1.794339457 |
| hsa-miR-888-5p | -0.248423363 | 1.589776113 |
| hsa-miR-889-3p | -0.333871954 | 1.633246384 |
| hsa-miR-891a-5p | -0.659162226 | 3.700347671 |
| hsa-miR-891b | -0.407085948 | 2.912082914 |
| hsa-miR-892a | -0.515982044 | 4.051579214 |
| hsa-miR-892b | -0.43509111 | 2.732880179 |
| hsa-miR-9-5p | -0.5425706 | 2.967529378 |
| hsa-miR-924 | -0.418454879 | 3.14481311 |
| hsa-miR-92a-1-5p | -0.434369599 | 1.696675567 |
| hsa-miR-92a-3p | -0.38141459 | 2.37567686 |
| hsa-miR-92b-3p | -0.806357344 | 3.387786079 |
| hsa-miR-933 | -0.908425964 | 5.82960703 |
| hsa-miR-934 | -0.279990538 | 1.62117063 |

| | | |
|---|---|---|
| hsa-miR-935 | -0.510761331 | 2.259594585 |
| hsa-miR-936 | -0.449698438 | 2.168451993 |
| hsa-miR-937-3p | -0.304172421 | 1.782376475 |
| hsa-miR-939-5p | -0.315965623 | 1.755911339 |
| hsa-miR-940 | -0.275777105 | 1.731941916 |
| hsa-miR-941 | -0.575934864 | 2.862881059 |
| hsa-miR-942-3p | -0.593338477 | 2.860401915 |
| hsa-miR-944 | -0.340877627 | 1.461110219 |
| hsa-miR-96-5p | -0.243472989 | 1.839772923 |
| hsa-miR-98-5p | -0.606569304 | 3.500349131 |
| hsa-miR-99a-5p | -0.53673085 | 2.376735767 |
| hsa-miR-99b-5p | -0.398202632 | 2.015559489 |
